# Conformational diversity in class C GPCR positive allosteric modulation

**DOI:** 10.1101/2023.11.07.565819

**Authors:** Giuseppe Cannone, Ludovic Berto, Fanny Malhaire, Gavin Ferguson, Aurelien Foullien, Stéphanie Balor, Joan Font-Ingles, Amadeu Llebaria, Cyril Goudet, Abhay Kotecha, Kutti R. Vinothkumar, Guillaume Lebon

## Abstract

The metabotropic glutamate receptors (mGlus) are class C G protein coupled receptors (GPCR) that form obligate dimers activated by the major excitatory neurotransmitter L-glutamate^1,2^. The architecture of mGlu receptor comprises an extracellular Venus-Fly Trap domain (VFT) connected to a transmembrane domain (7TM) through a Cysteine-Rich Domain (CRD). The binding of L-glutamate in the VFTs and subsequent conformational change results in the signal being transmitted to the 7TM inducing G-protein binding and activation ^3–6^. The mGlu receptors signal transduction can be allosterically potentiated by positive allosteric modulators (PAMs) binding to the 7TMs, which are of therapeutic interest in various neurological disorders^7–9^. Here, we report the cryoEM structures of metabotropic glutamate receptor 5 (mGlu_5_) purified with three chemically and pharmacologically distinct PAMs. We find that PAMs modulate the receptor equilibrium through their different binding modes, revealing how their interactions in the 7TMs impact the mGlu_5_ receptor conformational landscape and function. In addition, we identified a PAM-free but agonist-bound intermediate state that is stabilised by interactions mediated by intracellular loop 2. The activation of mGlu_5_ receptor is a multi-step sequential process in which the binding of the PAMs in the 7TM modulates the equilibrium towards the active state.

## Main

The major excitatory neurotransmitter in the mammalian nervous system, L-glutamate binds to various receptors in the membrane to modulate the activity at the synapse^7^. Among them, the metabotropic glutamate receptors (mGlu) are class C G-protein receptors. This family is composed of eight subtypes, divided into three groups based on their pharmacology, sequence similarity, and primary cellular signalling pathways. The structure of mGlu receptor is modular with an extracellular venus-fly trap domain (VFT) that binds the agonist L-glutamate and the cysteine-rich domain (CRD), which connects the VFT to the transmembrane domain (7TM)^2^. Typical of all class C GPCRs, mGlu receptors also form obligate homo- and heterodimers and oligomerization is essential for their biological function. Binding of the orthosteric agonist, L-glutamate induces the closure of the VFTs that mediates a large conformational change resulting in the signal being transmitted to the 7TM and binding of the G-protein^3–6,10,11^. Surprisingly, no natural ligands that bind to the 7TM of mGlu receptors have been identified yet but several small synthetic molecules can bind to 7TM, acting as positive or negative allosteric modulators, PAM and NAM, which enhance or inhibit receptor signalling, respectively.

The different mGlu receptors are widely expressed in the human brain where they modulate glutamatergic transmission, thus contributing to many important physiological processes^7^. Aberrations in glutamatergic transmission result in many neurological disorders, making mGlu receptor targets of therapeutic interest^7,8,12^. The mGlu_5_ subtype is an important modulator of synaptic plasticity involved in learning and memory. The mGlu_5_ positive allosteric modulation has strong potential for alleviating different symptoms of schizophrenia^9^. However, developing safe mGlu_5_ PAMs is a difficult task since some of them have been associated with adverse side-effects^13,14^. Understanding the effects of PAMs is a real challenge, and how they bind and impact on the dynamics of the mGlu activation mechanism is of great physiological and pharmacological interest.

Pharmacologically, PAMs trigger diverse signalling outcomes, resulting in reported biased signalling^15–17^ likely associated with multiple receptor conformations. Amongst the large number of PAMs that have been developed till date, some are defined as pure PAM that only potentiate the glutamate action (VU0409551^15^ and VU29^18^), but they can display a low agonist-like activity (intrinsic agonist activity) depending of the signalling pathway and cellular context^16,19–21^. Others PAMs, such as VU0424465^14^, are called ago-PAM that display a strong intrinsic agonist activity and can induce receptor signal transduction without the need for glutamate binding to the VFTs, either by activating the 7TM domain alone or by displacing the equilibrium between the inactive and active state of the receptor.

Several studies have reported the PAM-bound conformation of class C GPCRs^6,10,11,22^ and how the PAM and the G protein modulate the receptor conformation^3–5,23^. The class C GPCR activation mechanism is asymmetric i.e., with only one G protein being activated at a time by the receptor dimer. The 7TMs form asymmetric interface in the activated state both in the presence and the absence of G protein^4,5,24^. In mGlu_2_ receptor, only one ago-PAM was found bound to the G protein-coupled 7TM^4^ but in other class C receptors, such as the calcium receptor it has been observed that PAMs are bound to both protomers in the absence of G protein, although they adopt different binding pose or mode^24^. Recent single molecule-FRET analysis proposed a multi-state process including an agonist bound intermediate-active conformation between the inactive and active state^25^. mGlu activation mechanism is a dynamic, multi-step process and PAMs are likely to modulate this receptor state equilibrium, promoting the stabilisation of the active state. Despite the conserved domain architecture in mGlu, the differences in the 7TM sequence provides an opportunity to selectively modulate the mGlu’s activity in an isoform dependent manner and may lead to the discovery and design of new molecules with therapeutic potential and high level of selectivity. Thus, understanding the mechanism of PAM binding and the activation of the receptor is highly relevant. While structures of mGlu_5_ 7TM with NAM bound have been reported^11,26–28^, the binding mode of PAM in the mGlu_5_ 7TM and their impact on mGlu_5_ conformations has remained elusive.

Here, using cryogenic-sample electron microscopy (cryoEM), we report structures of full-length mGlu_5_ in detergent micelles bound to high affinity orthosteric agonist quisqualate and to chemically distinct PAMs: the ago-PAM VU0424465 and the PAMs VU0409551 and VU29. Combined with molecular pharmacology, we decipher the diversity of PAM binding mode and how they impact mGlu_5_ receptor conformation and function, providing insights into the effect of PAMs on the conformations and activity of mGlu_5_ receptor.

### Structures of PAM-bound mGlu_5_ receptor

The structures of PAM-bound mGlu_5_ receptor was solved by taking advantage of thermostabilised mGlu_5_ receptor that contains five thermostabilising mutations (T742A^5.42^, S753A^5.53^, T777A^6.42^, I799A^7.29^, A813L^7.43^) and truncation at its C-terminus after residue A856 (mGlu_5_-Δ856)^11,29^. We have previously shown that the thermostabilised receptor is fully functional, retaining its capability of activating the G protein when expressed in HEK293 cells^11,29^. The first structure described here is the ago-PAM VU0424465-bound mGlu_5_ receptor at a global resolution of 3.1 Å (we define ago-PAM VU0424465 as PAM1, data set 1 as PAM1_d1) with no symmetry imposed. Density is well defined in the ECD but is of lower resolution in the 7TM, a common feature in all the data sets described in this study (**Supplementary Fig. 1, 2**). The density for inner TM helices is better resolved and there is unambiguous density for the PAM in both protomers and for the surrounding residues (**Fig. 1A and B; Extended Data Fig. 1 and 7; Supplementary Fig. 1**). We were also able to build the loop ICL2 in this data set (using the deepEMhancer^30^ maps), which has been implicated in G-protein binding. A second independent data set of PAM1 bound receptor was collected with cold FEG and Falcon 4i detector housed at the bottom of Selectris energy filter and this led to obtaining of two different conformations of the receptor, at overall resolutions of 3.2 Å (PAM1, conformer 1 in dataset 2 as PAM1_d2_c1) and 3.5 Å (PAM1, conformer 2 in dataset 2 as PAM1_d2_c2) respectively (**Extended Data Fig. 2**). The difference between these two conformations includes some variability in PAM occupancy with global rmsd of 1.5 Å and 1.6 Å for the monomers. In these two conformations, the VFT (0.3 Å rmsd for Cα) overlay well but the CRD and the TMD show greater deviations (0.6 Å and 0.8 Å rmsd respectively). The PAM density is observed in both protomers but the conformer 1 has slightly weaker density when compared to PAM1_d2_c2, and PAM1_d1 (**Extended Data Fig. 7**). In addition, model superposition shows that these two conformations are the mirror of each other where PAM1_d2_c1 superimpose very well with PAM1_d1 (rmsd 0.7 Å and 0.8 Å for chain A and B respectively), while PAM1_d2_c2 (1.7 Å rmsd for chain A and 1.6 Å rmsd for chain B) differs from PAM1_d1 (**Fig. 2D and E**). These independent data sets provide confidence in the mode of PAM binding. We observe an asymmetrical interface previously described for other class C receptor that includes mGlu_2_^4^, CaSR^24^ and several mGlu heteromers^3^. All the structures with PAM bound in both protomers (PAM1_d1, PAM1_d2_c1, PAM1_d2_c2), has the top of TM6 shifted towards the extracellular side of the receptor (**Supplementary Fig. 3**). Altogether, this suggest that the mGlu_5_ receptor can adopt a minimum of two discrete conformations with both agonist and PAM bound, highlighting the dynamic nature of the mGlu_5_ dimer active state, prior to G protein binding.

**Fig.1.**
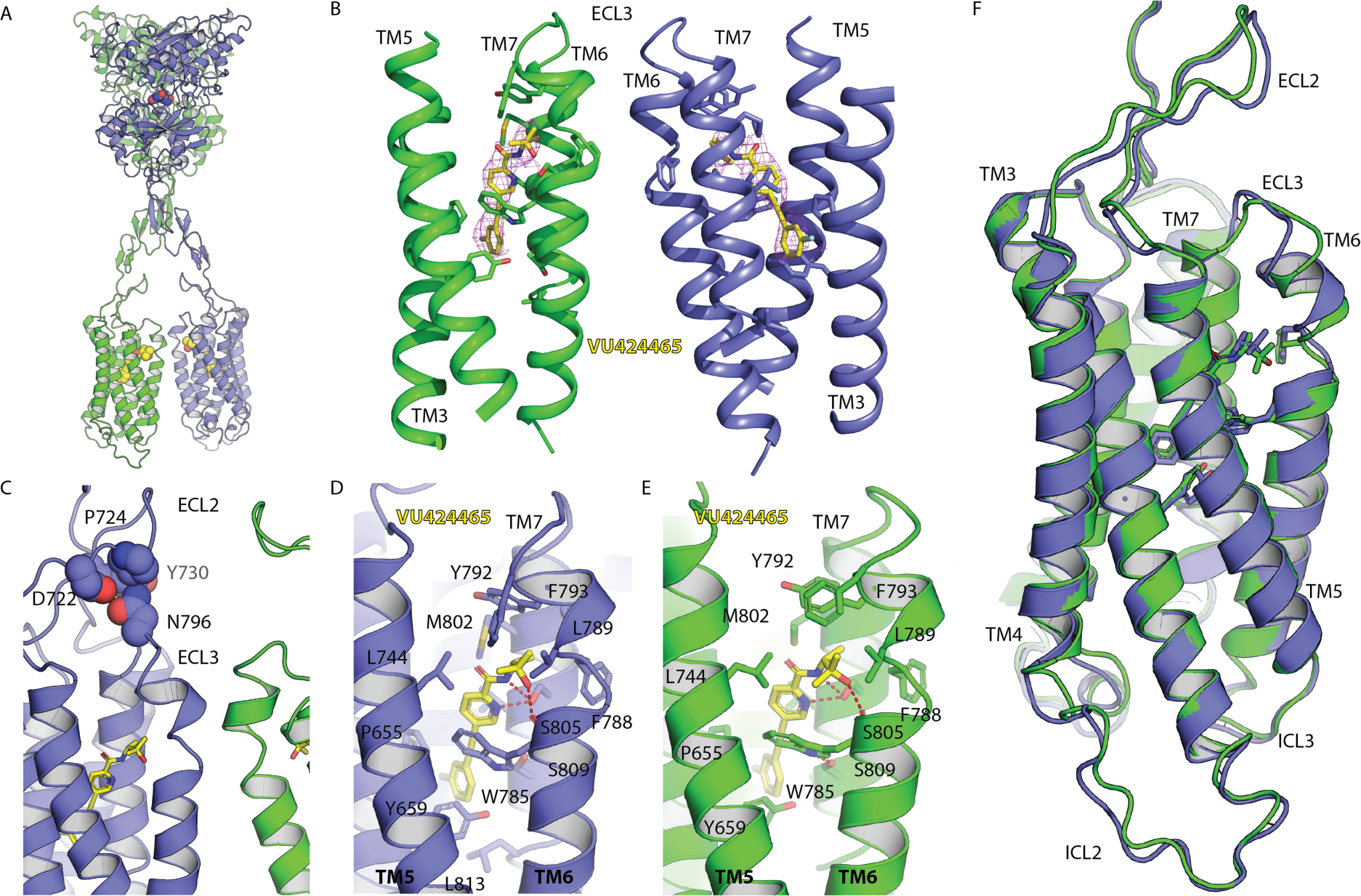
CryoEM structure of orthosteric agonist quisqualate and VU0424465 ago-PAM-bound conformation of the mGlu_5_ receptor. (A) mGlu_5_ structure display symmetrical binding of the ago-PAM VU0424465 to both 7TM as well as both VFTs display bound quisqualate. (B) CryoEM map (magenta) sharpened with B=-95 Å^2^ at 8 σ showing the density for the ago-PAM VU0424465 in both monomers surrounded by the TM helices and key side chain residues are shown in stick representation (carbon atoms in blue and green) and ligand carbon atoms in yellow. (C) The active state is stabilised by a limited number of residues localised at the top of TM6 from both 7TM, with an asymmetrical distribution. Molecular contact between N796 (ECL3) and D722, P724, Y730 (ECL2) observed only in one protomer. (D, E) Ligand binding site of VU0424465 in each protomer. VU0424465 make polar interaction with S805 and with the W785 carbonyl. It is stabilised by hydrophobic contact involving M802, V806, S809 in TM7; W785, Y793, F788, L789 in TM6 pushing out F788. (F) VU0424465 induces a shift of TM6 along its helical axis and towards the extra cellular side of the receptor. We could resolve the density for ICL2 that displays an extended helix 3 that stretches out of the detergent micelles. Figures were generated using pymol.

We then performed cryoEM reconstructions of the PAMs VU29 and VU0409551 at global resolutions of ∼3.2 Å (PAM2) and 3.0 Å (PAM3), respectively (**Extended Data Fig. 3 and 4**). The VU29 map had reasonably well-defined PAM density in only one of the monomers that allowed model building of the PAM (**Fig. 2A and Extended Data Fig. 7**) and of few bulky side-chains around the PAM binding sites. The second protomer displayed poor density for the ligand suggesting a lower occupancy and thus no PAM was built into this monomer in the map, but the residual density indicates a similar binding pose (**Extended Data Fig. 7**). In the case of VU0409551, only weak density for the PAM was observed in both protomers, preventing the model building (PAM3_c1) of this PAM. However, this data set has well resolved VFT allowing us to build few water molecules in both the protomers and the region around the agonist binding site and the 7TM is also better resolved than for VU29 structure (**Supplementary Fig. 2**).

**Fig.2.**
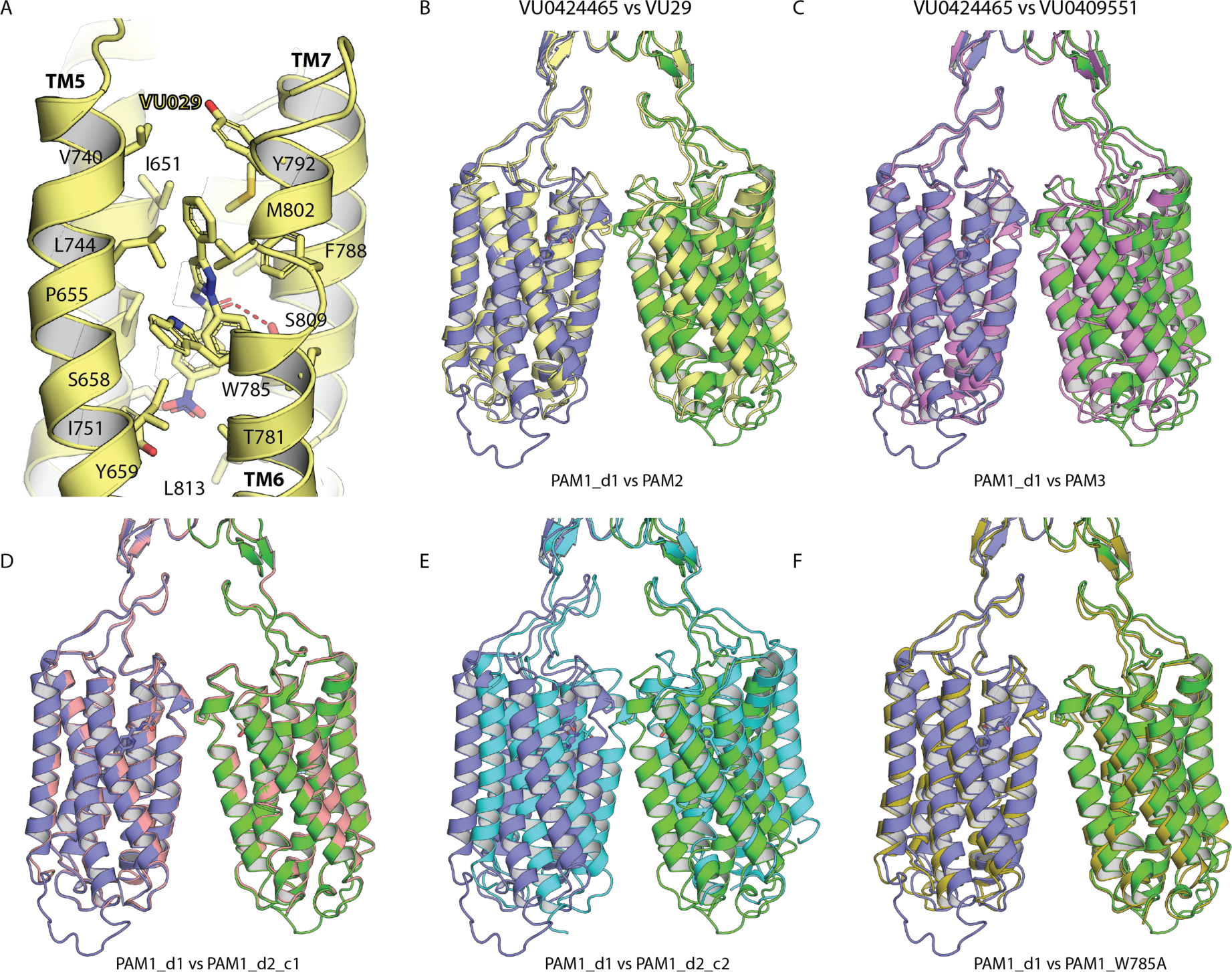
Conformational diversity of positive allosteric modulation of mGlu_5_ receptor dimer. (A) CryoEM structure of the PAM VU29-bound mGlu_5_ with only one molecule modelled and the VU29 binding mode. (B-F) Structure comparison of mGlu_5_ receptor dimer bound to ago-PAM VU0424465, PAMs VU29 and VU0409551 and VU0424465-bound to mGlu_5_ W785A mutant was performed. 3D alignment of residues 25-191 and 331-461 of both lobe-1 of the PAM1_d1 data set (one protomer in green and the other in blue) (VU0424465-bound mGlu_5_ receptor) was performed for VU29 (PAM2, yellow); VU0409551 (PAM3, light purple); the different conformations of VU0424465 including PAM1_d2_c1 (salmon) and PAM1_d2_c2 (cyan) and VU0424465-bound mGlu-5 W785A mutant (PAM3_c2, olive).

Subsequently, we also determined the structure of the thermostabilised receptor with an additional mutation, W785A (W785^6.50^), a key residue for NAM and PAM binding and signal transduction^31^. We co-purified the full-length mGlu_5_-5M W785A mutant bound to the ago-PAM VU0424465 and the 3D reconstruction has a global resolution of ∼3.4 Å (**Extended Data Fig. 5**). Removing the bulky W785 side chain strongly impairs the PAM occupancy but reveals some density found in the same region as other data sets (PAM_d1, PAM_d2_c1 and _PAM_d2_c2). The ligand is not modelled but the map implicates the role of W785 in VU0424465 ago-PAM binding.

When processing multiple PAM-bound data sets (3 different PAMs and one mutant), we also identified population of particles visually identical to mGlu_5_ inactive state receptor (i.e., with the TM domain separated apart) but only in the PAM3 data set, the number of particles was sufficient for medium-resolution EM reconstruction to ∼4.1 Å (**Fig. 3; PAM3_c2; Extended Data Fig. 6**). This structure reveals a closed conformation (c) of the VFTs, with quisqualate bound, but stabilised in an inactive (R) state, which we propose to be in an intermediate-active conformation (**Fig. 3**). The statistics of data processing and model building of all data sets are provided in Extended Data Table 1.

**Fig.3.**
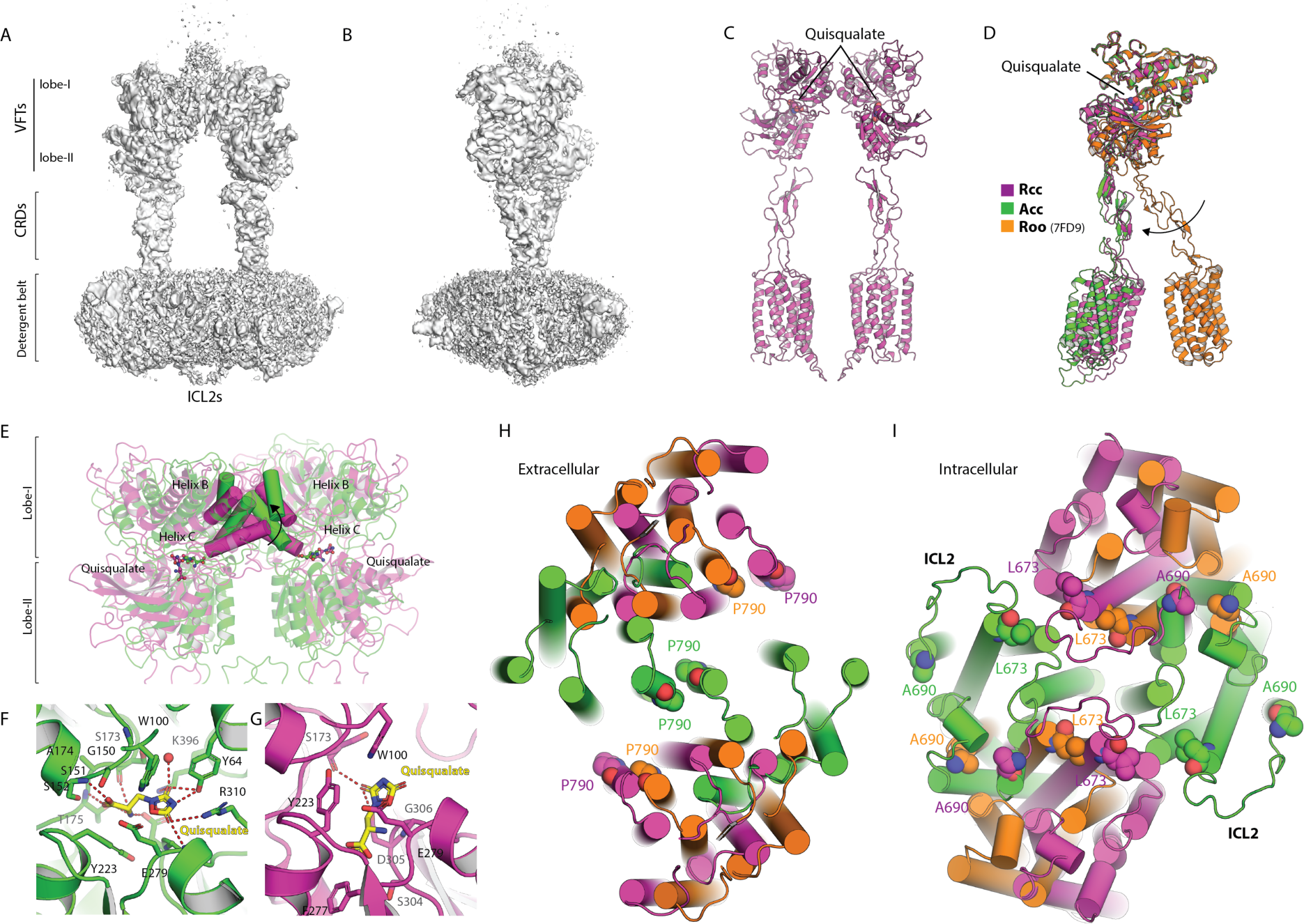
CryoEM structure of intermediate-active state (Rcc) with quisqualate bound but PAM free. (A and B) Two views of cryoEM map of the Rcc conformation illustrating molecular interaction between the ICL2 loops. (C) 3D model of the Rcc quisqualate-bound state show closed conformation of the VFTs induced by quisqualate binding. (D) Comparsion of monomers of Rcc model with antagonist-bound Roo state (PDB 7FD9) and the agonist and PAM-bound Acc state (PAM1_d1; green) conformation. (E) Superimposition of VU0424465-bound mGlu_5_ structure (PAM1_d1; green) and intermediate active state (Rcc) bound quisqualate but PAM free (PAM3_c2; purple). (F and G) Quisqualate standard binding pose as observed in VU0409551-bound mGlu_5_ structure and its alternate binding mode in intermediate-active state (Rcc). (H and I) The 7TMs of Rcc quisqualate-bound state undergo a tilt along the perpendicular axis to the membrane as P790 at the top of TM6 moves aways from each other compared to fully inactive state, before they come into close proximity in the Acc active state of the receptor. In the intracellular side the ICL2s cross and meet in the Rcc intermediate active state. ICL2 movement is illustrated by the L673 and A690 since the loop is not modelled in the fully inactivated state with open VFTs (o) and resting conformation (R), Roo (PDB 7FD9).

### mGlu_5_ PAM binding mode

Previously, the class C GPCR structures of mGlu_2_ revealed that only one ago-PAM is bound in the 7TM whereas two PAMs are bound in case of CaSR but in different binding modes ^4,24^. Nevertheless, despite the differences, an asymmetrical dimer interface was reported in both cases. In contrast, we observe ago-PAM bound in similar binding pose in both protomers in mGlu_5_ (**Fig.1B**).

VU0424465 (PAM1) shares a common binding site with the NAMs such as alloswitch-1^11^ or M-MPEP^28^ (**Supplementary_Fig. 4**). The fluorophenyl group is found between G624^2.45^, I625^2.46^, G628^2.49^, S654^3.39^, P655^3.40^, V806^7.36^, S809^7.39^ with one specific additional interaction with A810^7.40^ in mol **A (Fig. 1D and E**). Although the PAM is positioned in the same binding pocket as NAM, the main differences are localised at the top of TM6, where the ago-PAM induces the shift of TM6 towards the extracellular side with a kink at P790^6.55^ for only one protomer (mol B). As a consequence, the PAM slightly tilted upwards (toward the vertical axis of the 7TM) and makes an interaction with the S658^3.43^ and L813^7.43^, one of the thermostabilising mutation introduced in the receptor^11^ (**Fig. 1D**). Y659^3.44^ is displaced towards T781^6.46^ (**Fig. 1F**). In the second monomer (mol A), the W785^6.50^ is stabilised by a polar interaction with S809^7.39^ as found in the inactive state of the receptor suggesting that only one protomer is activated.

The pyridinecarboxamide of VU0424465 occupies the same pose in both protomer making van-der-walls interaction with M802^7.32^, L744^5.44^, and W785^6.50^ in TM6 (**Fig. 1D and E**). For both VU0424465 molecules, the N-pyridine atom makes polar contact to the S805^7.35^ and to the W785 carbonyl. The dimethylpropyl group of VU0424465 molecule contacts L744^5.44^, F788^6.53^, V789^6.54^, Y792^6.57^, F793^6.58^ in both protomers, whereas in mol B it is shifted towards TM7 and makes an additional interaction with T801^7.31^. Overall the PAM is slightly straightened in mol B, pushing TM6 up, towards the extracellular side.

The binding modes in PAM1_d2_c1 and PAM1_d2_c2 are very similar. Notably PAM1_d2_c1 model display a hydrogen bond between S809^7.39^ - W785^6.50^ in mol A only, similar to PAM1_d1. The PAM1_d2_c2 model displays the same polar contact in mol A, whereas the second protomer has an additional interaction between Y659 and T781^6.46^. The small differences observed could be because of the resolution and the quality of the maps in particular at the TMD (**Extended Data Fig. 1, 2**). With all these structures we can now define the signature of mGlu_5_ PAM bound state.

VU29 occupies a binding site similar to VU0424465 but with some significant differences. The main difference comes from W785^6.50^ being pushed away from the helical bundle by the phenyl ring of VU29 whereas F788 moves in to stabilise the PAM (**Fig 2A**). VU29’s nitrobenzamide group is localised in a similar position compared to the fluoro phenyl moiety of VU042465, but is likely to push away Y659^3.44^. VU29 is further stabilised by interactions with S658^3.43^ and a polar contact with S809^7.39^. In addition, VU29 engages with several residues from TM5 including V740^5.40^ and I751^5.51^. The PAM VU29 is the most affected by mutation of residues in the allosteric binding pocket (**Fig. 4C and D**). The 7TM has poor resolution and only few side-chains could be built with confidence, suggesting a dynamic nature of VU29 binding and, at the moment, we cannot exclude other binding modes.

**Fig.4.**
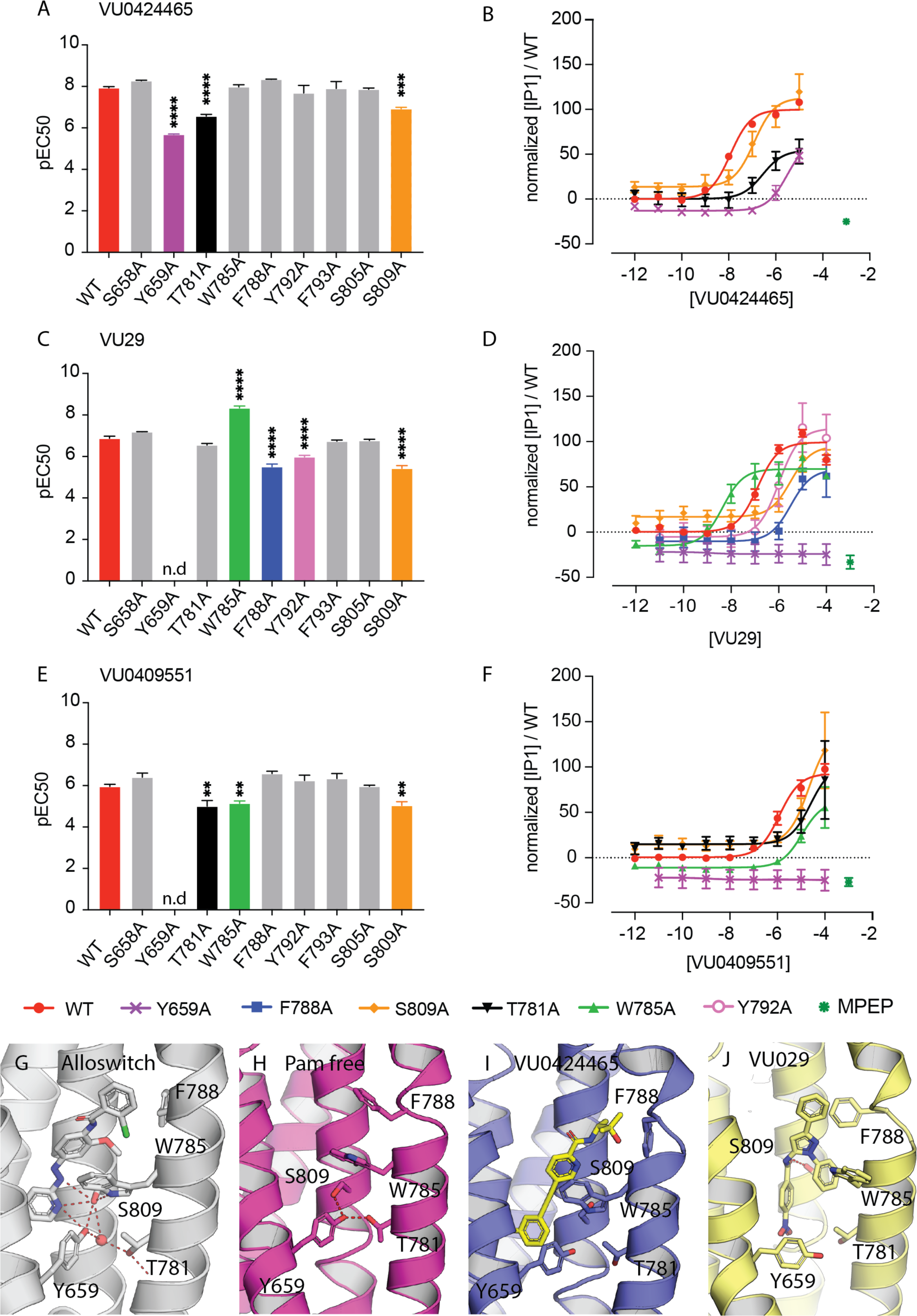
Functional role of PAM binding site residues on mGlu_5_ activity. We evaluated the influence of different residues of the PAM binding site in the 7TM of mGlu_5_ on the ability of the different PAM’s to potentiate agonist-induced activity. Single residues were mutated into alanine and functional consequences were determined by measuring the production of inositol monophosphate (IP_1_) in HEK293 cells transiently expressing the WT and mutated mGlu_5_ receptors. Experiments were performed in the presence of a fixed dose of the orthosteric agonist quisqualate (10 nM) and different doses of PAMs: VU0424465 (A and B), VU29 (C and D) and VU0409551 (E and F). Bar plots in A, C and E are pEC50 average values of at least 3 independent experiments +/−SEM. Dose response curve B, D and F are presented for mutants displaying statistically significant effect as compared to the WT type receptor. Each curve is an average of 3 independent experiments. Network of polar interaction between Y659, S809 and W785 is reframed by PAM binding (G-J). mGlu_5_ inactive state relies on W785, Y659, S809 forming polar contact with the NAM alloswitch-1 ligand (G; PDB 7P2L). W785 and F788 are both inwards orientated preventing the binding of the PAM in the 7TM PAM-free (H; Supplementary Fig. 4), Y659, S809 and W785 making polar contact as in NAM bound conformation. Ago-PAM VU0424465 binding destibilises polar interactions between Y659, S809 and W785 (I). VU29 pushes away Y659 and W785 but is stabilised by F788 (J). VU0424465 and VU29 stabilise different 7TM conformations.

### An asymmetric but flexible dimer interface

We have determined six 3D reconstructions for the PAM-bound mGlu_5_ receptor. It includes two conformations of the receptor bound to VU0424465, one bound to VU29, one to VU0409551, and one for the mGlu_5_-5M-W785A receptor bound to VU0424465. Surprisingly, only two of the VU0424465 structures (PAM1_d1 and PAM1_d2_c1) display an almost identical backbone superposition, from the VFTs to the 7TM when superposing the dimer (**Fig. 2**). When comparing monomers of each structure (i.e., PAM1_d1; **Fig1 F**), we observed that all VU0424465 structure (PAM1 models) display an asymmetrical interface, with a shift and a kink at P790 of TM6 for only one monomer, both much less pronounced in the W785 mutant and two other PAM structures (**Fig. 2 B, C and F; Supplementary_Fig. 3**). These are also the ones with the best occupancy for the PAM in each protomer. The 7TM interface is weaker for the two other PAMs suggesting that the lower occupancy in the binding site is likely associated to the weaker TM6-TM6 interface. This is also clearly reflected for the VU0409551 for which we were not able the model the PAM.

Even more surprising is the disruptive effect of the mutation W785A that affects not only the binding as reflected by the poor occupancy of VU0424465 but also disruption of the asymmetric interface between the 7TM as compared to PAM1 structure (PAM1_d1, d2_c1, d2_c2; **Fig. 2D, E, F**). Residue W785 is key for stabilising the VU0424465 ago-PAM in the allosteric binding site that mediates the active state asymmetrical dimer interface (**Fig. 1B, C D and F; Supplementary Fig. 3**). Indeed, in the VU29, VU0409551 and in the VU0424465+W785A mutant models, the distance between the Cα of P790 that is localised at the top of TM6 from each monomer significantly increase from ∼7.5 Å in the VU0424465 models (PAM1-d1, d2-c1 and d2-c2) up to 11 Å for the VU29 structure. Interestingly, VU0424465 has the highest affinity and is likely to better stabilise the active conformation, at least in the absence of the G protein and thus we conclude that the occupancy of the PAM in 7TM strongly affects the stability of 7TM dimer interface and propensity of the mGlu_5_ receptor active state.

### Intermediate active-state, a quisqualate-bound Rcc conformation

The class C GPCRs undergo a number of conformational changes both in their ECD as well in the 7TM. In absence of glutamate or presence of orthosteric antagonist, the VFTs adopt an open (o) state and a resting (R) conformation, with respect to lobe-I orientation^2,25^. Binding of L-glutamate or analogue such as quisqualate induces the closure of the VFTs and the reorientation of lobe-I that brings the lobe-II in proximity. This movement is translated to the 7TM that are far apart in the open and resting state of mGlu_5_, but form a dimeric interface in the active (A) state^6,11^. The multiple states that the receptors can adopt have been defined based on the conformation of VFT as open or close and the receptor as resting or active^32^. Thus, the antagonist bound receptors typically adopt Roo conformation^11^ (for the resting and open conformation of both VFTs) and the fully active receptor are called Acc (for active and closed conformation of both VFTs)^6,11^. The receptor can also adopt other conformations including the Rcc conformation as proposed for the mGlu_2_ receptor^25^.

Both the agonist and the PAM have been included throughout the purification of the receptor used in the study, and we still identified a sub-set of particles that showed distinguishable 2D classes compared to the PAM-bound active state in the data sets presented in this study. We could calculate only low-resolution maps from particles of the VU0424465 and VU29 data sets, but a reasonable medium resolution map at ∼4.1 Å was reconstructed from the PAM3 data set (**Extended Data Fig. 6**) and has slight anisotropy when compared to other data sets (**Supplementary Fig. 6**). The overall map of the receptor dimer is well resolved and the 7TMs are separated apart, as previously reported in the antagonist LY341495-bound mGlu_5_ receptor (**Fig. 3A, B, C and D**), although this structure represents an additional and different conformation of the receptor (**Fig. 3D, H and I**).

In the map, the VFT of one of the monomers is better resolved than the other and shows quisqualate bound in the classical pose as in other structures determined. The ligand is stabilised by several polar interactions (Y64, S152, S173, T175, G280, D305, R310, K396) and van-der-walls contact (W100, G150, S151, A174, Y223, E279, G306,) (**Fig. 3E, F, G**). In the second VFT, the quality of the map is lower indicating flexibility and the quisqualate adopts a different pose, the carboxylic group orients towards Lobe-II, making fewer van-der-walls contacts with residues Y233, E279, F277, S304, G306, and two polar interactions with D305 and S173, and suggesting an intermediate and transient binding mode (**Fig. 3G; Supplementary Fig. 5**). The weak density for one of the VFT is reflective of the receptor dynamics. Indeed, the 3D variability analysis (3DVA) of this subset of particles and subsequent analysis of the models by phenix.varref^33^ showed that one VFT is closed and in a stable conformation whereas the second VFT samples multiple states, from a closed to a more open conformation, compatible with quisqualate being weakly bound (**Supplementary Movie 1 and 2**). Although, we could not build the Lobe-1 in this monomer confidently due to poor density, we however can see that the helices B and C are starting to move and to undergo the transition from the R towards the A state when compared with the orthosteric antagonist LY341495-bound VFT conformation (**Supplementary Fig. 5**). This transition suggests that whilst one VFT is stabilised in the closed state, the second VFT is oscillating in multiple states between open (o) and close (c) conformations displacing the receptor equilibrium towards a more active-like A state initiating the motion to be translated to the CRD and to the 7TM. The results presented here illustrate the sequential but dynamic events occurring initially at the VFTs.

An interesting feature of this intermediate state is the ICL2, which extends from TM3, similar to the VU0424465-bound conformation (PAM1_d1), (**Fig. 3A, B, C; Extended Data Fig. 6**). In the Rcc conformation, the ICL2 from each protomer face each other, most likely stabilising the intermediate (**Fig. 3C and I**). From the intracellular side of the receptor the motion of TM3 clearly shows that ICL2 represents an intermediate state between the fully inactive state bound LY341485 and the active state presented here (**Fig. 3I**). The ICL2 loops of mGlu_5_ has a unique motif KKICTKK and mutating any of these residues affects the efficacy of the PAM signalling with a dramatic effect in particular for the residue I680 (**Extended Data Fig. 8; Supplementary Table 1**). The ICL2 loop has been proposed to be important for G protein interaction of class C GPCRs^34^ and it is tempting to speculate that this intermediate conformation prevents the binding of the G-protein.

Despite a modest global resolution for this state of the receptor, several key residues involved in PAM binding have defined density but no clear density could be detected for the PAM. The side-chains of W785^6.50^ and F788^6.53^ are pointing towards the helical bundle (**Fig 4H**). Although W785 stabilises the VU04244645 in PAM1 models (**Fig. 1**), the presence of F788 is likely to cause steric hindrance and allows us to exclude the possibility that the PAM in the 7TM have lower occupancy in the intermediate active state Rcc. We defined this conformation as a PAM-free intermediate-active state. On hindsight, this observation supports the view that the PAMs stabilise the active Acc conformation. The weaker the PAM, the less stabilised is the Acc state with agonist quisqualate bound in the VFT, allowing particles to distribute between the intermediate-active Rcc and the Acc state. Ago-PAM that induces substantial mGlu_5_ receptor signalling in absence of orthosteric agonist is likely due to the propensity of the PAM to stabilise the mGlu_5_ receptor active state, although we cannot exclude that PAM can activate isolate 7TMs^35^ i.e., even in the Rcc conformation^11^.

### mGlu_5_ PAM activation mechanism

The selection of the PAMs for this study was initially based on their difference in chemical scaffold, affinity, as well as their different intrinsic agonist activity. In the literature, VU0424465 is described as an high affinity ago-PAM^14^ contrary to VU29 and VU0409551, which are less able to activate mGlu_5_ by themselves ^16,20^. Here, we confirm that VU0424465 possesses a high intrinsic agonist activity in absence of the agonist quisqualate, being able to almost fully activate mGlu_5_ at 1 µM (83 ± 6 % of the maximal activation induced by quisqualate in our assay) (**Extended Data Fig. 9; Supplementary Table 2**). In contrast, VU29 and VU0409551 solely display a low intrinsic agonist activity, being able to partially activate the receptor at 1µM (20 ± 9% and 15 ± 1% of maximal activation induced by the agonist quisqualate for VU29 and VU0409551, respectively) (**Extended Data Fig. 9; Supplementary Table 2**).

W785^6.50^, a key residue for GPCR activation mechanism^31^, differentially affects the activity of the PAMs (**Fig. 4; Supplementary Table 3**). The mutation of this residue to alanine has no significant effect on the signalling activity induced by VU0424465 (**Fig. 4A and B**), likely due to its strong affinity. In contrast, the potency of the other two PAMs VU29 (**Fig. 4C and D**) and VU0409551 (**Fig. 4E and F**) is significantly affected in the W785A mutant (**Supplementary Table 3**). The allosteric effect of VU29 is improved in this mutant but the effect by VU0409551 is reduced. In the model proposed here, VU29 binding pushes the W785^6.50^ side chain out and thus removing the side-chain is likely to facilitate its efficient binding to the receptor (**Fig. 2A**). In the other words, when mutating W785^6.50^ in alanine, more space is created for the PAM VU29 than for ago-PAM VU0424465 and VU0409551.

Another key residue for all three PAM is Y659^3.44^. Replacing this tyrosine abolishes the VU29 and VU0409551 modulation and strongly impairs the effect of VU0424465, although some activation was measured (**Fig. 4; Supplementary Table 3**). Y659 is localised at the bottom of the binding site and was found in a network of polar contact in the inactive conformation of the 7TM structure solved to high resolution by X-ray crystallography^11^ (**Fig. 4G**). Here, the Y659^3.44^ is displaced towards T781^6.46^ in PAM1_d1 and moves away from S809 disrupting the Y659^3.44^ - S809^7.39^ interaction. The Y659^3.44^ also interacts with T781^6.46^ in one monomer only for the model PAM1_d2_c2. Pharmacology presented here highlight the role Y659^3.44^ but also of T781^6.46^, that display reduced potency for VU0424465 as well as for VU0409551. In the protomer where the VU29 is modelled, the nitrobenzamide group pushes the Y659^3.44^ towards TM5 and the interaction with S809 is also broken.

A third important residue is the F788 that undergoes the most dramatic movement when all structures are compared (**Fig. 4G-J**). In the Rcc conformation, the side chain of F788^6.53^ is facing the core of 7TM (**Fig. 4H**) as previously reported for the M-MPEP bound conformation, and is likely to prevent the PAM binding (**Supplementary Fig. 4**). The position of the F788^6.53^ may adopt multiple conformations without affecting the pharmacology of PAM or NAM. Indeed, in the structure of alloswitch-1, a photoswitchable NAM has forced the F788^6.53^ out (**Fig. 4G**) but maintains strong hydrogen bond network with S809^7.39^, Y659^3.44^ and W785^6.50^. However, in presence of PAM VU0424465, F788^6.53^ is facing out (**Fig. 4I**), whereas it moves and interact with VU29, stabilising its binding as suggested by its reduced potency in F788A mutation (**Fig. 4C, D and J**).

We also observe interactions between N796 (ECL3) and D722, P726 and Y730 (ECL2) of the active-like protomer in the PAM1_d1 model that may help to stabilise the active conformation (**Extended Data Fig. 10; Supplementary Table 4**). Such interaction is also observed in the two other VU0424465 structures (PAM1_d2_c1 and PAM1_d2_c2). However, mutating these residues has only weak effect on the efficacy of PAM.

The structural and functional analysis of ago-PAM and PAM-bound receptor presented here reveals a dynamic/flexible network of contacts between residues in the ligand binding site stabilising the mGlu_5_ receptor active state and to ultimately promote G protein binding. The global differences of the 7TM between the inactive state and the active state are not extensive and revolve around the residues Y659^3.44^, W785^6.50^, S809^7.39^, F788^6.53^ and T781^6.46^. These residues have been shown to be important for receptor inactive state and for transmitting receptor signalling^36^. The analysis of the pharmacological characterisation clearly illustrates that residues Y659^3.44^, S809^7.39^, and T781^6.46^ affect PAM’s signal transduction, as an activation module. Associated with the structural characterisation presented here, the W785^6.50^ appears to provide account for the complexity of mGlu_5_ allosteric modulation and for the diversity of PAM binding mode.

## Discussion

Allosteric modulation of class C GPCRs is the subject of intense research with the grail of discovering high affinity and selective molecules for specific receptor subtype. The main therapeutic potential for mGlu_5_ PAMs is in the treatment of different symptoms of schizophrenia^9^. However, the development of a safe mGlu_5_ PAM with antipsychotic properties has remained a real challenge^14,15^. One of the main reasons is the complexity of the mGlu_5_ signal transduction and the adverse side effects associated with some of these PAMs in preclinical studies. The complex signalling of mGlu_5_ receptor has been associated to biased signalling^17^. Conformational diversity of the class C GPCRs is also directly linked to signalling profiles of the receptor but remains elusive.

Here, we selected three different PAMs with different scaffold and signalling profile. The VU0424465 is defined as an ago-PAM whereas as VU29 and VU0409551 are considered as pure PAM for biased signalling as observed in specific cellular context^16,37^. They also display different affinity that translates into different potency ^18,14,15^. In fact, VU0424465 displays the highest affinity and potency, while VU0409551 has the weakest potency of the PAM, as analysed by the Gq protein activation and inositol phosphate-1 production. (**Fig. 4; Supplementary Table 3**).

The high occupancy and better density of the ago-PAM VU0424465 reported in this study correlates with higher affinity than PAM VU29^18^ and VU0409551^14^. The receptor occupancy affects the conformation and stability of the mGlu_5_ active state (**Fig. 2B and C**), correlating with the difference in receptor signal transduction in cells i.e., a higher proportion of mGlu_5_ receptor in the active state will bind and activate the G protein. An important observation is that the ago-PAM VU0424465 is bound to both protomers and stabilises the asymmetric interface characteristic of class C receptor dimer and better stabilises the active state (**Fig 1 and 2**). We observe density in both protomer for the PAMs VU29 and VU0409551 albeit at lower occupancy, which probably allows the receptor to explore more conformations.

Perhaps due to the lower affinity of PAM3, a larger population of the receptor is able to exist in the closed VFT and in the inactive resting (R) state allowing us to determine the structure of an active intermediate state (Rcc) conformation. The L-glutamate analogue, quisqualate, is bound to both VFTs but adopts different poses: a standard pose, as previously reported^6,11^ and seen in all active state solved here, and an alternate pose that is likely to be energetically unfavourable, suggesting significant degree of VFT dynamics (**Fig. 3E, F and G, Supplementary movies 1 and 2**). This analysis strongly suggests a sequential conformational change that starts with the binding of quisqualate, or L-glutamate, and the stable closure of one of the VFTs that will promote the closure of the second one. However, this first step brings the two ICL2s at proximity that consequently stabilises the Rcc state, hampering the movement of the 7TM that likely requires the stable closure of the second VFT and the reorientation of Lobe-I from R to A state in order for the receptor to reach the fully active state. The molecular interaction between ICL2 identified in this intermediate-active state might transiently stabilise the receptor in the Rcc state and keeps the 7TM apart (**Fig. 3C and I**). Additionally, the ICL2 intermolecular contact likely blocks the binding of G protein, making this conformation an intermediate state that needs to be overcome before mGlu_5_ reaches the active state and G protein binding. In summary, ICL2s of the Rcc state may act as a checkpoint in preventing activation of the mGlu_5_ receptor in absence of L-glutamate in the *in-vivo* context. Orthosteric agonist quisqualate favours the formation of this Rcc conformation, increases the probability of the receptor to overcome this intermediate conformation and to reach the active state. Once both VFTs are stabilised in the Acc state, the 7TMs are in close proximity and the PAM will provide further stabilisation to the active state.

We have unveiled the binding site for the ago-PAM VU0424465 and identified a binding pose for VU29 illustrating the differential contribution of TM6 residues mainly W785^6.50^ and F788^6.53^ **Fig (4I and J**). Functional analysis confirms the importance of the mGlu_5_ receptor residues W785^6.50^, F788^6.53^, Y659^3.44^, S809^7.39^ and T781^6.46^ in PAM allosteric modulation, which is highly valuable in the design of new PAMs.

Comparing the PAM-free and PAM-bound conformations allowed us to illustrate the motion of key residues that control the binding of PAM and subsequently stabilise the active state, by sampling the 7TM conformation landscape. PAMs differentially displace the Y659^3.44^ and F788^6.53^. The W785^6.50^ is central to differential stabilisation of the PAM in the binding site, either positively for VU0424465 and VU0409551 or negatively for VU29. Mutating this residue to alanine has a direct effect on both the PAM occupancy and the active state conformation that may be comparable to low affinity PAM such as VU0409551. The differences between the inactive and active state at the 7TM is subtle and in agreement with the absence of TM6 motion as defined as the molecular signature for the class A and B GPCR active conformations^38^. Indeed, comparison of the structures of NAM-bound mGlu_5_ show some similarity between the NAM and PAM binding site (**Supplementary Fig. 4**)^26,28^. The residues Y659^3.44^, S809^7.39^, W785^6.50^ are making tight polar network that also involves T781^6.46^ through a water molecule, in which S809 was shown important for mediating NAM effect^11^. Here we show that this tight network of polar interaction is reorganised, and key interaction such as Y659^3.44^-S809^7.39^ polar interaction is destabilised. Y659^3.44^ is a key residue for PAM signal transduction and for NAM inhibition of the receptor. Interestingly, Y659A mutant has no effect on signal transduction induced by the orthosteric agonist quisqualate, suggesting that the receptor dimer can still sample the 7TM asymmetric dimer interface, in absence of PAM. The strong stabilising potential of the ago-PAM VU0424465 on the receptor active state is likely to trigger a much higher level of signal transduction than orthosteric L-glutamate, and may induce possible undesired side-effect.

The multi-step activation process of the mGlu_5_ receptor perhaps includes internal check-points such as the ICL2 and the conformations of the residues in the PAM binding sites that can be modulated by small molecules and biologics. The strong stabilising potency of an ago-PAM such as VU0424465 is likely to trigger a much higher level of signal transduction by potentiating orthosteric L-glutamate as would a pure PAM perform, but also due to intrinsic agonist activity that triggers signal transduction in absence of L-glutamate, likely giving account for undesired side-effect. Thus, the structures reported here provide a template for exploring positive allosteric modulation and to be integrated in structure-based drug discovery program for identifying new PAM for mGlu_5_ receptor.

## Methods

### Expression and purification of the human full-length thermostabilized mGlu_5_ receptor bound to activators

The thermostabilised mGlu_5_-5M receptor and mGlu_5_-5M_W785A mutants were expressed and purified as previously reported^11^. mGlu_5_-5M receptor and mGlu_5_-5M_W785A constructs were subcloned into the BacMam vector. The constructs contain peptide signal from baculovirus GP64 envelope surface glycoprotein of the *Autographa californica* nuclear polyhedrosis virus (AcNPV baculovirus), the flag (DYKDDDDK), 10xHis tag, and the precision protease recognition site (LEVLFQGP) at the N terminus, followed by the human mGlu_5_-5M or mGlu_5_-5M_W785A. The construct also contains a H350L mutation, initially designed for binding of a nanobody and a N445A mutation to remove a glycosylation site. The mGlu_5_-5M_W785A includes the additional mutation W785A and the N445A glycosylation mutation was reversed.

High-titre recombinant baculovirus was obtained using the Bac-to-Bac Baculovirus expression system (Invitrogen) as described by the manufacturer. Briefly, Sf9 cells grown at 28°C in EX-CELL 420 medium (Sigma Aldrich) were infected at a density of 3×10^6^ cells using human mGlu_5_-5M and mGlu_5_-5M_W785A virus and P2 virus were harvested by centrifugation 48 hours post-infection. HEK293S cell line lacking N-acetylglucosaminyltransferase I^39^ (HEK293S GnTI^−^) cells were grown at 37°C in humidified 5% CO_2_ in shaking incubator (110 rpm/minute) in Freestyle medium complemented by 10% of Fetal Bovine Serum (Sigma Aldrich). HEK293S GnTI (-) cells were infected at a density of 3×10^6^ cells per mL with 6% of freshly produced P2 virus (60 mL per litre) and 10 mM sodium butyrate was added 24 hours post-infection. Cells expressing mGlu_5_-5M and mGlu_5_-5M_W785A were harvested 96 hours post-infection and stored at −80°C for further purification.

For membrane preparation, frozen cell were resupended in a lysis buffer containing 25 mM HEPES (pH 7.4), 10 mM MgCl_2_, 20 mM KCl, 5 mM EDTA, 1 mM PMSF (Sigma Aldrich), and the complete protease inhibitor cocktail tablet (Roche). The pellets were washed two times with the lysis buffer and an additional wash performed using high salt concentration of sodium chloride (25 mM HEPES (pH 7.4), 10 mM MgCl_2_, 20 mM KCl, 1 M NaCl). Membrane fraction was harvested by centrifugation at 45,000 rpm (≈200.000*g*) for an hour and resuspended with a lysis buffer supplemented with 40% glycerol and then stored at −80°C. Membrane preparation from 2 litres of cell culture expression were resuspended into a final volume of 120 mL of 25 mM HEPES buffer (pH 7.4), 0.4 M NaCl, 10% Glycerol,10 µM quisqualate (Abcam), 10 mM iodoacetamide (Sigma), and the complete protease inhibitor cocktail tablets and complemented with 10 µM of either PAM such as, VU0424465^11^, PAMs VU29 (med chem express) VU0409551 (R&D system), The mixture was then incubated at room temperature (RT) for 1 hour and the membranes were solubilised by adding 1% (w/v) n-Dodecyl-β-D-Maltopyranoside (DDM, Anatrace), and 0.2% (w/v) cholesteryl hemisuccinate (CHS, Anatrace) for 1.5 hours at 4°C, under shaking. Solubilised receptors were harvested by centrifugation at 45,000 rpm for 1 hour and supplemented with 10 mM imidazole (Sigma) before loading onto a Nickel-NTA (Ni^2+^) resin (HisTrap HP, GE Healthcare) at 0.5 mL/min, overnight at 4°C. The resin was pre-equilibrated in 25 mM HEPES (pH 7.4), 400 mM NaCl, 10 µM quisqualate, 10 µM of PAM, 5% (v/v) glycerol, 0.2% (w/v) DDM, 0.04% (w/v) CHS, and 10 mM imidazole. The resin was washed using a step gradient of imidazole with concentrations of 10mM, 20mM, 40mM and 50 mM and the receptor was eluted with 25 mM HEPES (pH 7.4), 400 mM NaCl, 10 µM quisqualate, 10 µM of PAM, 5% (v/v) glycerol, 0.05% (w/v) DDM, 0.01% (w/v) CHS, and 250 mM imidazole. The eluted receptors were incubated onto a Flag affinity resin (Anti-Flag M1, Sigma) for 1 hour at 4°C, washed with 10 column volumes of a wash buffer (25 mM HEPES (pH 7.4), 150 mM NaCl, 10 µM quisqualate, 10 µM PAM, 0.03% (w/v) DDM, 0.006% (w/v) CHS, and 2 mM Ca^2+^). The receptor was eluted with 0.2 mg/mL Flag peptide in a buffer containing 25 mM HEPES pH 7.4, 0.15 M NaCl, 10 µM quisqualate, 10 µM VU0424465, 0.03% DDM (w/v), 0.006% CHS (w/v), 2 mM EDTA. The eluted fractions were then concentrated using a Amicon Ultra-4 centrifugal concentrator 100 kDa (Millipore), centrifuged for 10 minutes at 80,000 rpm to eliminate aggregates, and then loaded on a size exclusion chromatography (SEC) column (Superdex 200 Increase 10/300, GE Healthcare) pre-equilibrated with 25 mM HEPES pH 7.4, 0.15 M NaCl, 10 µM quisqualate and 10 µM PAM, 0.03% DDM (w/v) and 0.006% CHS (w/v). Eluted and purified mGlu_5_-5M receptor and mGlu_5_-5M_W785A receptor mutant bound to quisqualate and PAM were concentrated to 5-6 mg/ml using Amicon Ultra-4 centrifugal concentrator 100 kDa (Millipore) for cryoEM grid preparation and analysis.

### CryoEM sample preparation, data collection and image processing

#### VU0424465 (PAM1) with K3 detector

The agonist quisqualate and PAM VU0424465 bound mGlu_5_ FL receptor in detergent micelles, at 5 mg/mL was applied to a Quantifoil^®^ Au 0.6/1.0 µm 300 mesh grids. Grids were rendered hydrophilic by glow-discharging for 10 seconds at 25 mA using a PELCO EasiGlow instrument. Grids were plunge-frozen using EM GP2 Plunge Freezer (Leica) with the chamber equilibrated at 95% RH and 4°C. Grids were blotted with Whatman no. 1 filter paper for 4.5-6 seconds and plunge frozen in liquid ethane. The frozen grids were stored in liquid nitrogen until further use. Data were collected on Titan Krios G3i (ThermoFisher) at the European Synchrotron Radiation Facility (ESRF, Grenoble, France) with a K3 detector in counting mode with a quantum energy filter and 20 eV slit width. Data was collected from one grid with a dose of ∼15.2 e^−^/p/s, and 2.4 second exposure at nominal magnification of 105000x, corresponding to pixel size of 0.84 Å. A total of 19830 movies were used for image processing. Movie files were imported into cryosparc 3.2^40^. Motioncorr was carried out with cryosparc in-built patch motion correction (multi) and CTF estimation was performed in-built patch CTF estimation (multi). Initial particles picking was performed using Blob picker in cryosparc and particles were initially extracted with large box size in order to have the particles centered and for subsequent analysis they were down sampled. From initial large set of particles obtained after 2D classification, *ab-initio* model generation was performed followed by several round of 2D and 3D classifications to select good particles and obtain final map using refined and reconstructed using non-uniform refinement^41^ (NU) with C1 symmetry. The complete workflow is described in Extended Data Fig. 1. Local resolutions were estimated with Relion^42^.

#### VU0424465 (PAM1) with Falcon4i detector

Another independent data set of the mGlu_5_ bound with quisqualate and PAM VU0424465 was collected with a Falcon4i detector (PAM1_d2). Grids were made by application of the receptor (6mg/ml) to a Quantifoil^®^ Au 0.6/1 µm 300 mesh grids. Grids were treated with 9:1 Ar:O_2_ plasma (in a Fischione 1070 chamber) for 60 s to render them hydrophilic. Grids were plunge-frozen using Vitrobot Mk IV (ThermoFisher) with the chamber equilibrated at 100% RH and 10°C. Grids were blotted with Whatman no. 1 filter paper for 5 seconds with a blot force of 10 without any waiting or draining time and plunge frozen in liquid ethane.

The data acquisition was carried out using the cold field emission gun (CFEG) Titan Krios G4 microscope (ThermoFisher), at the ThermoFisher NanoPort facility in Eindhoven. The CFEG microscope was equipped with a Falcon 4i detector operating in electron counting mode and was integrated at the base of a Selectris Energy Filter. The microscope featured fringe-free imaging (FFI) and aberration-free image shift (AFIS)^45^, enabling the acquisition of multiple images before stage movement.

Data was collected with a sampling rate of 0.727 Å per pixel, employing an electron flux of ∼5 e^−^/Å²/s and a total fluence of 50 e^−^/Å². Movies were processed using Relion 4.0^42^. A total of 19,115 movie frames underwent motion correction using Relion’s built-in program, followed by contrast transfer function (CTF) estimation carried out with CTFFIND4^43^. Initial particle picking was executed using Topaz with a specified threshold of −2^44^. Reference-free two-dimensional (2D) classification was employed to identify high-quality particles. A subset of these particles was then employed to train a new Topaz model, which was subsequently utilized for repicking particles. These particles were extracted with a box size of 360 pixels and down sampled to speed up processing. From the 2D classification, a total of 396,207 particles were selected and subsequent 3D classification resulted in 221,875 particles. These selected particles were refined in 3D with C1 symmetry, resulting in a resolution of 3.5 Å after postprocessing, as determined by Fourier shell correlation (FSC) at 0.143. The map was further enhanced through one round of Bayesian Polishing and two rounds of CTF refinements following the method outlined by Zivanov et al. (2018)^45^, resulting in a final resolution of 2.7 Å. C2 symmetry expansion and 3D classification with focus on the TMD (using a T value of 128 and no image alignment) revealed the presence of two conformers. Particles corresponding to these two conformers were re-refined separately in Relion^42^.

Particles associated with these two conformers were subsequently imported into Cryosparc v4.3.0^40^. Duplicate particles arising from symmetry expansion were eliminated before the import. Non-uniform (NU) refinement, local CTF refinement, and local refinement were carried out independently for each conformer. These refinements yielded overall resolutions of 3.2 Å (2.9 Å) and 3.5 Å (3.0 Å) for conformer 1 (PAM1_d2_c1) and conformer 2 (PAM1_d2_c2), respectively (the numbers in brackets are the resolution with tight mask and corrected FSC in cryosparc). The complete workflow is described in Extended Data Fig. 2. Local resolutions were estimated with Relion^42^.

#### VU29-bound mGlu_5_ receptor

The agonist quisqualate-bound mGlu_5_ WT in detergent micelles, at 5 mg/mL and PAM VU29 was applied to a Quantifoil^®^ Au 0.6/1.0 µm 300 mesh grids. Grids were rendered hydrophilic by glow-discharging for 10 seconds at 25 mA using a PELCO easiGlow instrument. Grids were plunge-frozen using EM GP2 Plunge Freezer (Leica) with the chamber equilibrated at 95% RH and 4°C. Grids were blotted with Whatman no. 1 filter paper for 4.5-6 seconds with no waiting/drying time and plunge frozen in liquid ethane.

The data acquisition was carried out using the cold field emission gun (CFEG) Titan Krios G4 microscope (ThermoFisher), at the ThermoFisher NanoPort facility in Eindhoven equipped with a Falcon 4i detector integrated at the base of a Selectris Energy Filter. Data was collected using fringe-free imaging (FFI) and aberration-free image shift (AFIS) with a sampling rate of 0.727 Å per pixel, employing an electron flux of ∼6.1 e^−^/Å²/s and a total fluence of 49.62 e^−^/Å². The processing of this data set was done with Cryosparc^40^. Movie files (9469) were imported into cryosparc 3.2. Motioncorr was carried out with cryosparc in-built patch motion correction (multi) and CTF estimation was performed in-built patch CTF estimation (multi). Initial particles picking was performed using Blob picker in cryosparc and particles were initially extracted with large box size in order to have the particles centered and for subsequent analysis they were down sampled. From initial large set of particles obtained after 2D classification, *ab-initio* model generation was performed followed by several round of 2D and 3D classifications to select good particles and obtain final map using refined and reconstructed using non-uniform refinement^41^ (NU) with C1 symmetry. The complete workflow is described in Extended Data Fig. 3. Local resolutions were estimated with Relion^42^.

#### VU0409551-bound mGlu_5_ receptor

The agonist quisqualate-bound mGlu_5_ WT in detergent micelles, at 5 mg/mL and PAM VU0409551 was applied to a Quantifoil^®^ Au 0.6/1.0 µm 300 mesh grids. Grids were rendered hydrophilic by glow-discharging for 10 seconds at 25 mA using a PELCO easiGlow instrument. Grids were plunge-frozen using EM GP2 Plunge Freezer (Leica) with the chamber equilibrated at 95% RH and 4°C. Grids were blotted with Whatman no. 1 filter paper for 4.5-6 seconds with no waiting/drying time and plunge frozen in liquid ethane. Data was collected on a Titan Krios G3i (ThermoFisher) at the European Synchrotron Radiation Facility (ESRF, Grenoble, France) with a K3 detector in counting mode with a quantum energy filter and 20 eV slit width. Data collection was performed with a dose of ∼18.4 e^−^/p/s, an exposure of 1.9 seconds at 105000x, corresponding to pixel size of 0.84 Å. A total of 19943 movies were used for image processing. Movie files were imported into cryosparc 3.2^40^. Motioncorr was carried out with cryosparc in-built patch motion correction (multi) and CTF estimation was performed in-built patch CTF estimation (multi). Initial particles picking was performed using Blob picker in cryosparc and particles were initially extracted with large box size in order to have the particles centered and for subsequent analysis they were down sampled for 2D classification. From initial large set of particles obtained by 2D classification, *ab-initio* model generation was performed, which identified two different conformations followed by several round of 2D and 3D classifications to select good particles and obtain final maps using using non-uniform refinement^41^ (NU) with C1 symmetry for both the active and the RCC conformation. For the Rcc conformation, 3D variability analysis in cryosparc was performed and the resulting maps was used for analysis with Phenix.varref^33^ (described below). The complete workflow are described in Extended Data Fig. 4 and 6. Local resolutions were estimated with Relion^42^.

#### mGlu_5_ W785A

The agonist bound mGlu_5_ W785A in detergent micelles, at 6 mg/mL with quisqualate and PAM VU0424465 was applied to a Quantifoil^®^ Au 1.2/1.3 µm 300 mesh grids. Grids were rendered hydrophilic by glow-discharging for 1 min at 25 mA using a PELCO EasiGlow instrument. Grids were plunge-frozen using Vitrobot Mk IV (ThermoFisher) with the chamber equilibrated at 100% RH and 16°C. Grids were blotted with Whatman no. 1 filter paper for 3-3.5 seconds with a blot force of 0 and plunge frozen in liquid ethane. The frozen grids were stored in liquid nitrogen until further use.

Data were collected on Titan Krios G3i (ThermoFisher) at the National CryoEM facility in Bangalore with a Falcon 3 detector in counting mode and faster data acquisition with AFIS^46^. Data were collected from two independent grids made on different days with a dose of ∼0.54 e^−^/p/s and an exposure of 60 seconds at 75000x, corresponding to pixel size of 1.07 Å. Images from both data sets were manually curated to remove crystalline and thicker ice, followed by import of movie files into Relion 4.0^42,45^. Motioncorr was carried out with relion’s in-built program and CTF estimation was performed with CTFFIND4^43^. Based on CTF resolution and upper limit of defocus to 3.5 μ, a total of 2751 and 3001 images from each batch were selected further processing (the complete workflow is described in Extended Data Fig. 5). Low-pass filtered templates from prior data sets were used for automated particle picking with Gautomatch (https://www2.mrc-lmb.cam.ac.uk/download/gautomatch-053/) and particles were extracted with 4.28 Å sampling and 2 rounds of 2D classification was performed to select good particles. These selected particles were extracted with a box size of 384 and 1.07 Å sampling and further enriched for good particles with 2D classification. The particles from batch 1 and batch 2 were individually refined in 3D with C1 symmetry resulting in 4.8 and 6.1 Å respectively after postprocessing (all resolutions described here are of FSC at 0.143). Subsequently, the particles were subjected to Bayesian Polishing^45^ and both B-factor weighted particles (shiny.star) were joined and refined. The combined data with 260,251 particles after 3D refinement and postprocessing resulted in a 4.3 Å map.

The combined data was subsequently imported into Cryosparc v4.3.0^40^ and ab-initio reconstruction was performed with 5 classes and maximum resolution of 12 Å followed by heterogenous refinement. Three of the best classes or volumes were combined (that had the same resolution) and both homogenous and Non-uniform (NU) refinement^41^ were performed resulting in 4.3 Å and 3.6 Å maps respectively. Local and global CTF refine was then carried out on the particles with the volume from NU refinement and subsequent refinement resulted in an overall resolution of 3.4 Å. Local resolution was estimated in relion and the local resolution of the maps revealed that the VFT of the receptor had higher resolution than the CRD and the TMD as observed for the other data sets described in this study.

For all the data sets, the anisotropy or the sphericity was estimated using the 3DFSC server^47^ (**Supplementary Fig. 6**).

### Model building and refinement

Previous agonist bound active state model (7fd8) was used for rigid body fitting into PAM1_d1 map using Chimera^48^ and manually inspected and rebuilt where required using Coot^49^. Multiple maps were used during model building in all data sets including the maps with different B-factor sharpening and the deepEMhancer maps^30^. For instance, the ICL2 loop in PAM1_d1 was built using the deepEMhancer map generated with mask option and for modelling water molecules, sharpened maps with automatic B-factor estimation were used. Ligand restraint files were generated using JLigand^50^. Omit maps were generated using servalcat^51^ and used to verify the ligands and solvent.

For the subsequent data sets, the refined model from PAM1_d1 was used as template and monomers were individually fit using Chimera. This global rigid body gave good fit for the VFT and the CRD and 7TM regions were manually fit to the density with Coot first using rigid body and then with real space refinement (in the case of W785A mutant, the residue was changed to alanine). Unlike the other data sets, in W785A mutant data set, there was also extra density at N445 for glycosylation and N445 was restored from alanine, but sugar has not been modelled. In all the data sets, the density for ICL2 loop is ordered to varying degree but only in PAM1_d1 and PAM3_c2, they have been modelled. For the PAM3_c2 data set, the VFT and CRD from PAM1_d1 was used and the 7TM was rigid body with 7fd9 as template along with 7p2L.

Water molecules in the VFT region of PAM1_d1 and PAM3_c1 were conservatively picked using Coot and they were manually checked using the hydrogen bonding distance and density as the criteria. Models were refined using Phenix^52,53^ in particular the real space refinement (ref) and Servalcat/Refmac in ccpEM^51,54,55^. For the real space refinement of the model with Phenix, the map sharpened map with half the value of B-factor estimated by the relion postprocess was used and when refmac was used for refinement, the half-maps were provided. The statistics of the final models are provided in Extended Data Table 1.

For the RCC conformation, the maps from 3D variability analysis and the model without any ligands was used as input to Phenix.varref^33^ to obtain the dynamic nature of the VFT (as shown in Supplementary Movies).

### Cell culture, transfections, expression and inositol phosphate one (IP_1_) accumulation assay

#### Cell culture and transfection

HEK293 cells (ATCC CRL-1573) were cultured in Dulbecco’s modified Eagle’s medium (DMEM) supplemented with 10% FBS and were maintained at 37°C in a humidified atmosphere with 5% CO_2_. Absence of mycoplasma was checked regularly to guarantee the use of mycoplasma-free cells. An alanine mutant library of the SNAP-tagged human truncated mGlu_5_ receptor (mGlu_5_-Δ856, designated as WT in the following experiments) was generated using Quick-change strategy (Agilent technologies) and verified by sequencing (Eurofins Genomics)^11^. Cells were transiently transfected using electroporation with the indicated mGlu_5_ construct. The DNA mixture for transient transfection included 0.6 µg of the indicated mGlu_5_ mutants, 2 µg of the glutamate transporter EAAC1 cDNA (to reduce the influence of glutamate that may remain in the assay medium as released by the cells) per 10 million of cells. Cells are seeded at the density of 100,000 cells per well into white bottom 96-well culture plates (Greiner Bio-one). Following 24 hours of transfection, expression and IP_1_ production are measured.

#### Cell-based pharmacological assays

Wild-type and mutant mGlu_5_ receptor expression was determined by labelling the SNAP-tag receptors with BG-lumi4-Tb. In brief, cells were incubated for 1 hour at 37°C with 100 nM of BG-Lumi4-Tb in 50 µL of glutamate-free DMEM GlutaMAX-I. Cells were then washed twice with Tag-Lite buffer, and then 100 µL of Tag-Lite buffer was added to each well (Revvity). Lumi4-Tb fluorescence was then measured with BMG Pherastar.

Before the commencement of the IP_1_ accumulation assay, HEK293 cells were incubated for 1.5 hours with glutamate-free DMEM GlutaMAX-I (Life Technologies), to reduce the extracellular concentration of glutamate. The IP_1_ accumulation assay kit (Revvity, France) was used for the direct quantitative measurement of IP_1_ in HEK293 cells transiently transfected with mGlu_5_ constructs. Cells were stimulated with various concentrations of allosteric compounds with a fixed concentration of the orthosteric ligand Quisqualate (10 nM) for 30 minutes at 37°C, 5% CO_2_. For the Schild experiments, cells were stimulated with various concentrations of Quisqualate with a fixed concentration of the allosteric compounds. Then, cells were lysed using the lysis buffer for 15min and half of the lysate was incubated with conjugate-lysis buffer containing the d2-labeled IP_1_ analog and the terbium cryptate-labeled anti-IP_1_ antibody in 384well plate, according to the manufacturer’s instructions. After 1 hour incubation at RT, the HTRF measurement was performed after excitation at 337 nm with 50 µs delay, terbium cryptate fluorescence and tr-FRET signals were measured at 620 nm and 665 nm, respectively, using a PheraStar fluorimeter (BMG Labtech).

Data were fitted by a 3 parameters equation and statistical analyses were perfomed using one-way ANOVA followed by a Dunnett’s post-hoc test, using GraphPad Prism version 10 (San Diego, CA).

## Data availability

The cryoEM density map and model coordinate have been depositied in the Electron Microscopy Data Bank (EMDB) and the Protein Data Bank (PDB) respectively, under the accession code : EMDB-37973 and PDB-8X0B (PAM1_d1, VU0424465- and quisqualate -bound mGlu_5_-5M, active state) ; EMDB-37974 and PDB-8X0C (PAM1_d2_c1, VU0424465- and quisqualate -bound mGlu_5_-5M, active state) ; EMDB-37975 and PDB-8X0D (PAM1_d2_c2, VU0424465- and quisqualate -bound mGlu_5_-5M, active state) ; EMDB-37976 and PDB-8X0E (PAM1, VU0424465- and quisqualate-bound mGlu_5_-5M-W785A, active state) ; EMDB-37977 and PDB-8X0F (PAM2, VU29- and quisqualate-bound mGlu_5_-5M, active state); EMDB-37978 and PDB-8X0G (PAM3_c1, VU0409551- and quisqualate-bound mGlu_5_-5M, active state); EMDB-37979 and PDB-8X0H (PAM3_c2, quisqualate-bound mGlu_5_-5M, intermediate-active state, Rcc).

## Acknowledgement

We acknowledge the MRC Laboratory Molecular Biology EM and computing core facilities (Cambridge, UK), the European Synchrotron Radiation Facility for provision of beam time on CM01 and we would like to thank Alessandro Grinzato and Gregory Effantin for assistance (Grenoble, France). We acknowledge the METi imaging facility, member of the national infrastructure France-BioImaging supported by the French National Research Agency (ANR-10-INBS-04). We acknowledge the platform of pharmacology Arpeges of the Institut de Génomique Fonctionnelle (Montpellier, France). We acknowledge the National cryoEM Facility at Bangalore for data collection. L.B. was supported by Ligue Contre le Cancer. GF was supported by SWITCH-ON (ANR-20-CE11-0019). J.F.-I., and A.L. were supported by Ministerio de Ciencia e Innovación, Agencia Estatal de Investigación and ERDF - A way of making Europe (PID2020-120499RB-I00) and by the Catalan government (2021 SGR 00508). G.C. was supported by Medical Research Council grant MC-U105184322. K.R.V. acknowledges the DBT B-Life grant DBT/PR12422/MED/31/287/2014, and the support of the Department of Atomic Energy, Government of India, Government of India, under Project Identification No. RTI4006. KRV is part of EMBO Global Investigator Network. G.L. was supported by SWITCH-ON (ANR-20-CE11-0019) and Ligue Contre le Cancer.

## Author contributions

GC and LB contributed equally to the work and share the co-first authorship. LB performed expression and purification of the full-length mGlu_5_-5M bound to quisqualate and three different PAM samples for cryoEM analysis, with the help of GF. LB, AF performed grid freezing. GC prepared cryoEM grids, screened, collected and processed cryoEM data. AK performed data collection at TFS cryoEM facility. FM performed and analysed pharmacological experiments on the mGlu_5_ receptor and receptor mutants under the supervision CG. JFI synthesized the VU0424465 under the supevision of AL. KRV and GL performed image processing and building/refinement of full-length mGlu_5_-5M models. KRV and GL interpreted the data and wrote the manuscript with contributions from all the authors. GL designed, supervised and coordinated the project.

---

**Extended Data Figure 1.**
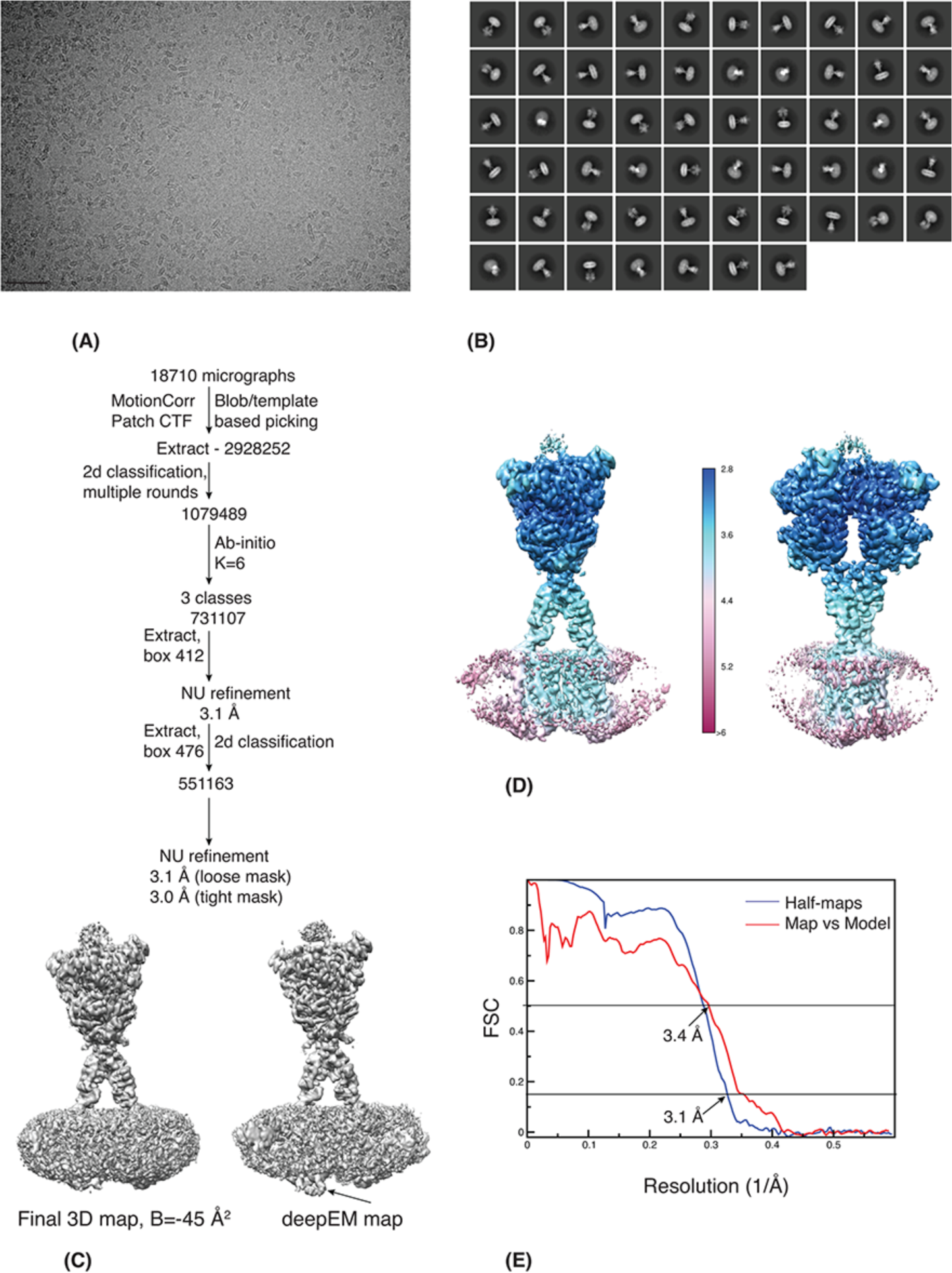
CryoEM analysis of VU0424465 and quisqualate bound to full-length mGlu_5_(mGlu_5_-Δ856) (A) A representative micrograph of the VU0424465 and quisqualate bound FL mGlu_5_ in detergent micelles imaged on ice and collected on K3 detector. Scale bar is 500 Å. (B) Select reference-free 2D class averages of mGlu_5_ with a box size of 476 pixels and sampled at 0.84 Å. (C) The image processing workflow of the mGlu_5_ with all the steps performed in cryosparc. The final 3D maps with sharpened with a B-factor of −45 Å^2^ and generated with deepEMhancer using the half-maps and a mask are shown. The arrow marks the ICL2 loop in the deepEM map that was used for model building. D) CryoEM maps coloured according to the local resolution plot of FL mGlu_5_ as determined with Relion 4.0 are shown in two different views. Much of the VFT is at a higher resolution and the CRD and the TMD are at lower resolution along with the detergent-lipid micelle. (E) The Fourier shell correlation (FSC) curves of two half-maps with a mask (blue) and the map and model (red) are shown. The estimated resolution by comparison of the two half-maps from refinement and postprocessing at FSC 0.143 is 3.1 Å and the one of map vs model at FSC 0.5 is 3.4 Å.

**Extended Data Figure 2.**
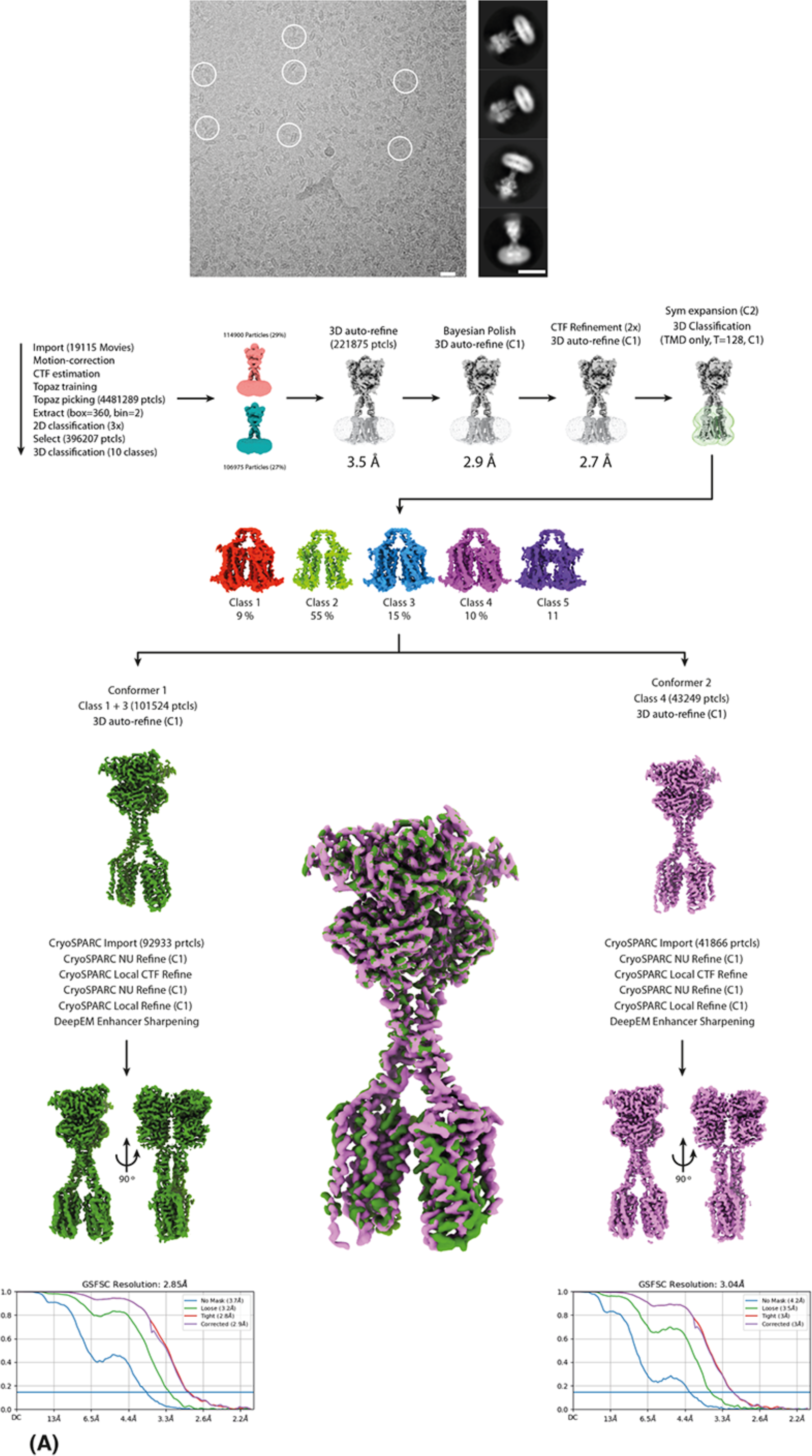
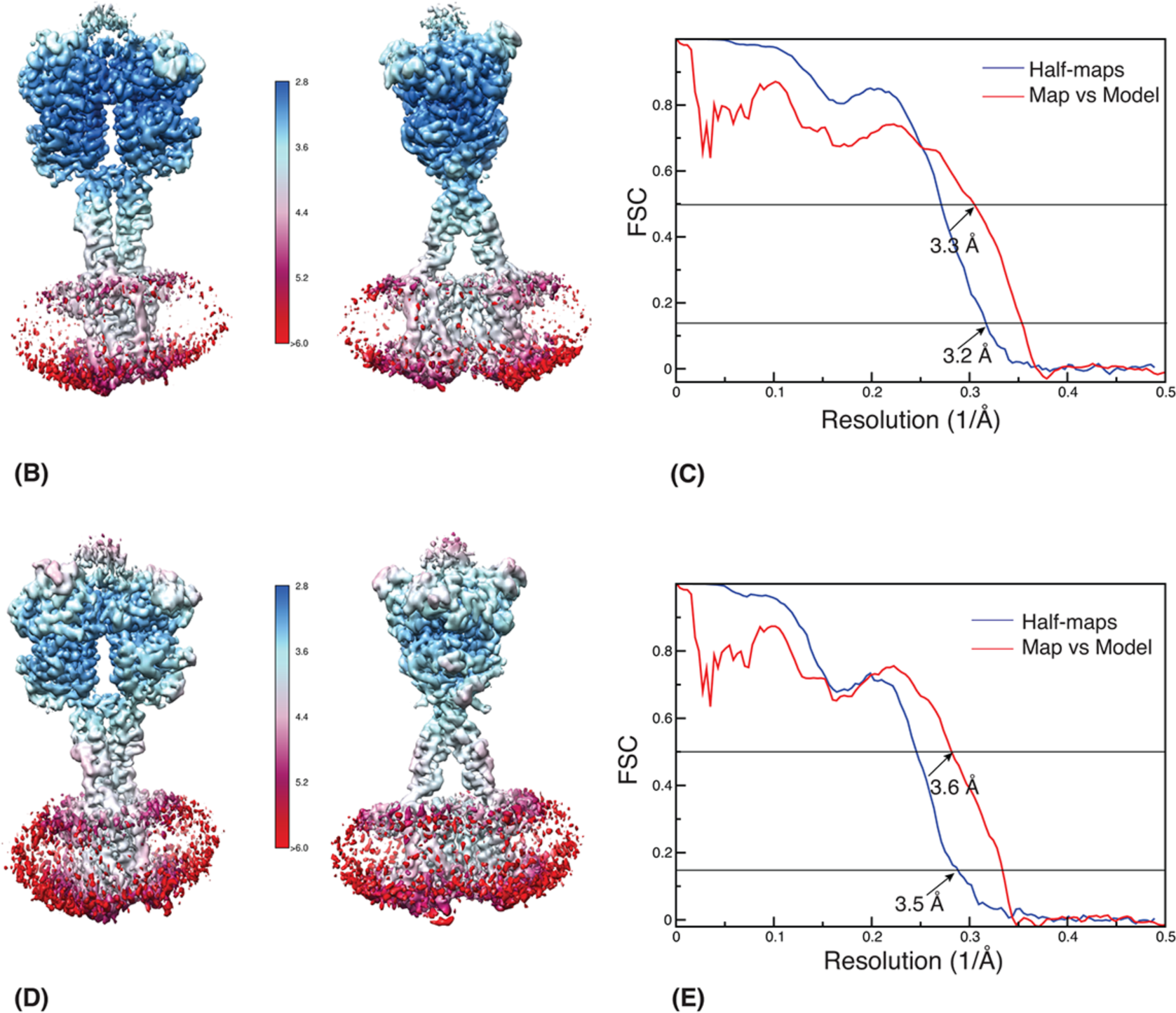
CryoEM analysis of VU0424465 and quisqualate bound to full-length mGlu_5-_5M (mGlu_5_-Δ856) with the data collected with Falcon 4i. (A) A representative micrograph of the VU0424465 and quisqualate bound FL mGlu_5_ in detergent micelles imaged on ice and collected on cFEG, selectris and Falcon IV detector, with some particles marked with white circle. Scale bar is 100 Å. Selected reference-free 2D class averages of mGlu_5_ with a box size of 256 pixels. The image processing workflow of the mGlu_5_ with the steps performed in Relion and cryosparc. B) CryoEM maps coloured according to the local resolution plot of FL mGlu_5_ conformer 1 as determined with Relion 4.0 are shown in two different views. Much of the VFT is at a higher resolution and the CRD and the TMD are at lower resolution along with the detergent-lipid micelle. (C) The Fourier shell correlation (FSC) curves of two half-maps with a mask (blue) and the map and model (red) are shown for conformer 1. The estimated resolution by comparison of the two half-maps from refinement and postprocessing at FSC 0.143 is 3.2 Å and the one of map vs model at FSC 0.5 is 3.3 Å. D) CryoEM maps coloured according to the local resolution plot of full-length mGlu_5_ conformer 2 as determined with Relion 4.0 are shown in two different views. Much of the VFT is at a higher resolution and the CRD and the TMD are at lower resolution along with the detergent-lipid micelle. (C) The Fourier shell correlation (FSC) curves of two half-maps with a mask (blue) and the map and model (red) are shown for conformer 2. The estimated resolution by comparison of the two half-maps from refinement and postprocessing at FSC 0.143 is 3.5 Å and the one of map vs model at FSC 0.5 is 3.6 Å.

**Extended Data Figure 3.**
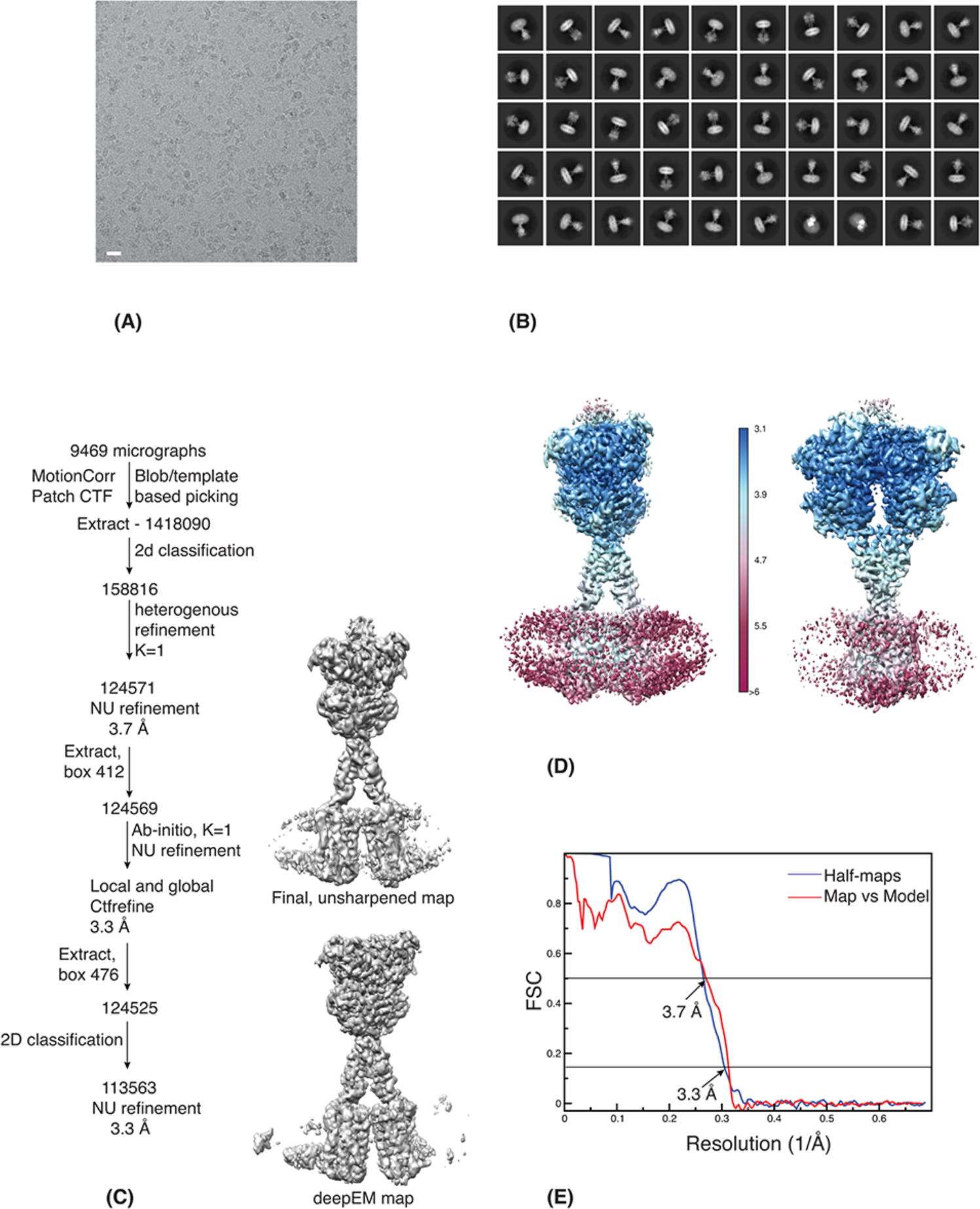
CryoEM analysis of VU29 and quisqualate bound to full-length mGlu_5_ (mGlu_5_-Δ856) (A) A representative micrograph of the FL mGlu_5_ with VU29 in detergent micelles imaged on ice and collected on Falcon 4i detector. Scale bar is 100 Å. (B) Selected reference-free 2D class averages of mGlu_5_ with VU29 and quisqualate bound with a box size of 476 pixels and sampled at 0.84 Å. (C) The image processing workflow of the mGlu_5_ with all the steps performed in cryosparc. The final 3D maps with the unsharpened and one generated with deepEMhancer using the half-maps and a mask are shown. D) CryoEM maps colored according to the local resolution plot of FL mGlu_5_ as determined with Relion 4.0 are shown in two different views. Much of the VFT is at a higher resolution and the CRD and the TMD are at lower resolution along with the detergent-lipid micelle. The TMD of the VU29 data set is of lower quality than the other data sets with PAM bound. (D) The Fourier shell correlation (FSC) curves of two half-maps with a mask (blue) and the map and model (red) are shown. The estimated resolution by comparison of the two half-maps from refinement and postprocessing at FSC 0.143 is 3.3 Å and the one of map vs model at FSC 0.5 is 3.7 Å.

**Extended Data Figure 4.**
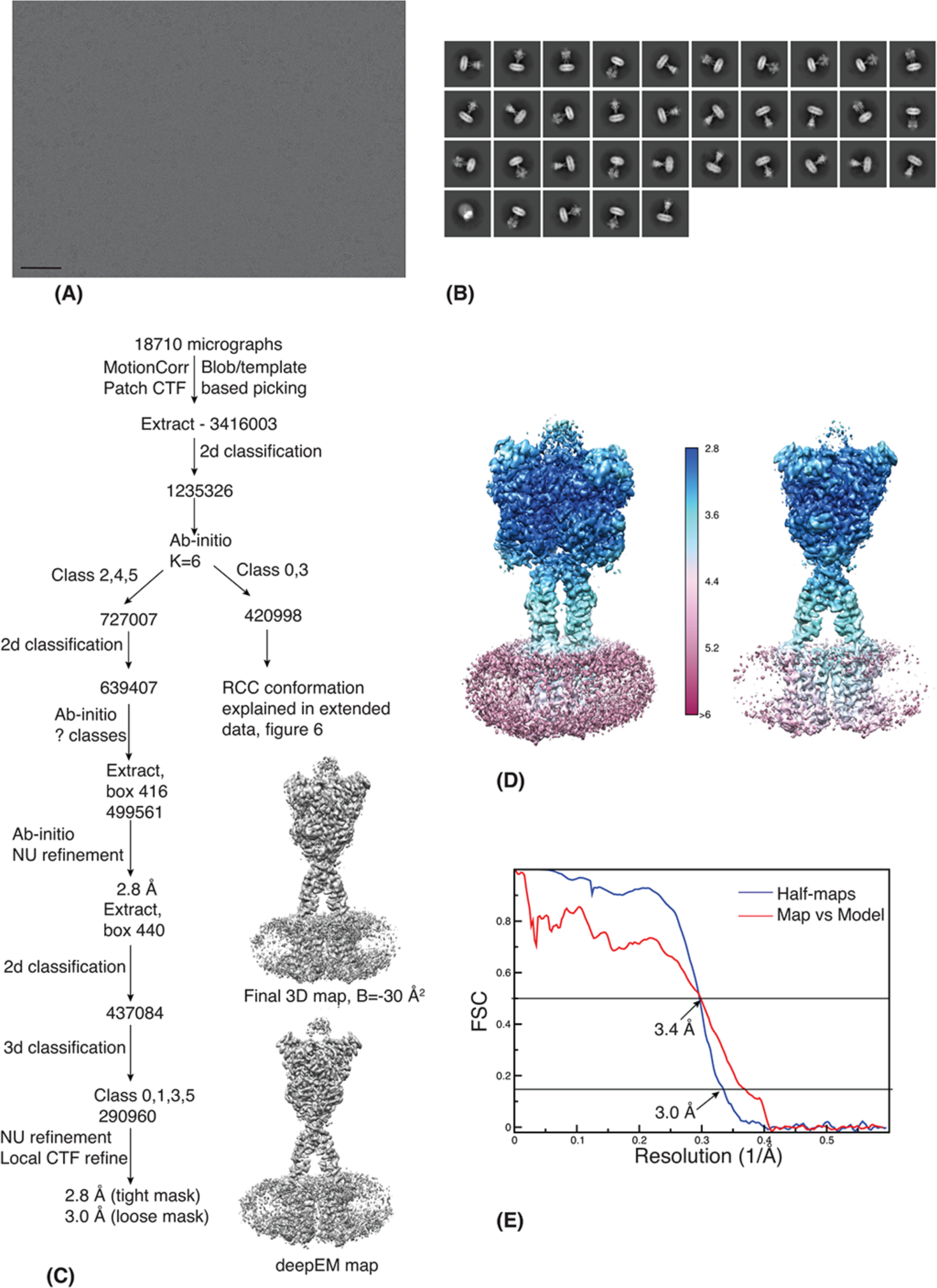
CryoEM analysis of VU0409551 and quisqualate bound to full-length mGlu_5_ (mGlu_5_-Δ856) in an active conformation. (A) A representative micrograph of the VU0409551 and quisqualate full-lenght mGlu_5_-5M in detergent micelles imaged on ice and collected on K3 detector. Scale bar is 500 Å. (B) Selected reference-free 2D class averages of VU0409551 mGlu_5_ with a box size of 440 pixels and sampled at 0.84 Å. (C) The image processing workflow of the mGlu_5_ with all the steps performed in cryosparc. The final 3D maps with either sharpened with a B-factor of −30 Å^2^ and generated with deepEMhancer using the half-maps are shown. D) Cryo EM maps coloured according to the local resolution plot of FL mGlu_5_ as determined with Relion 4.0 are shown in two different views. Much of the VFT is at a higher resolution and the CRD and the TMD are at lower resolution along with the detergent-lipid micelle. (D) The Fourier shell correlation (FSC) curves of two half-maps with a mask (blue) and the map and model (red) are shown. The estimated resolution by comparison of the two half-maps from refinement and postprocessing at FSC 0.143 is 3.0 Å (the tight mask in cryosparc estimates 2.8 Å) and the one of map vs model at FSC 0.5 is 3.4 Å.

**Extended Data Figure 5.**
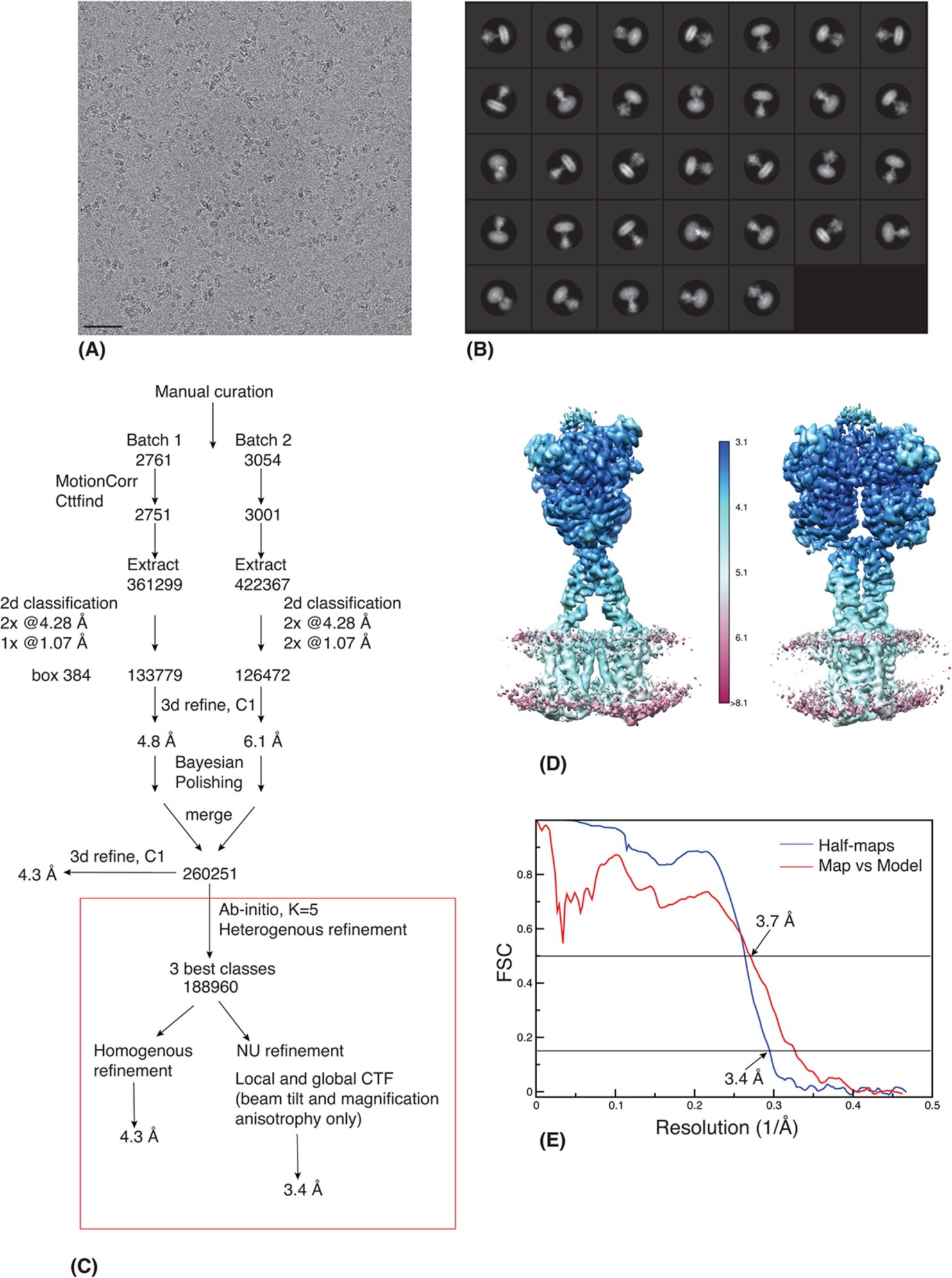
CryoEM analysis of VU0424465 and quisqualate bound mGlu_5_ W785A mutant. (A) A representative micrograph of the full-length mGlu_5_-5M_W785A in detergent micelles imaged on ice and collected on Falcon 3 detector. Scale bar is 500 Å. (B) Selected reference-free 2D class averages of mGlu_5_ W785A mutant with a box size of 384 pixels and sampled at 1.07 Å. (C) The image processing workflow of the mGlu_5_ W785A mutant. All the initial steps in image processing were performed with Relion 4.0 and the steps shown in red rectangle were performed with cryosparc 4.3.0. D) CryoEM maps coloured according to the local resolution plot of FL mGlu_5_ W785A mutant as determined with Relion 4.0 are shown in two different views. Much of the VFT is at a higher resolution and the CRD and the TMD are at lower resolution along with the detergent-lipid micelle. (E) The Fourier shell correlation (FSC) curves of two half-maps with a mask (blue) and the map and model (red) are shown. The estimated resolution by comparison of the two half-maps from refinement and postprocessing at FSC 0.143 is 3.4 Å (the tight mask after correction in cryosparc estimates 3.3 Å) and the one of map vs model at FSC 0.5 is 3.7 Å.

**Extended Data Figure 6.**
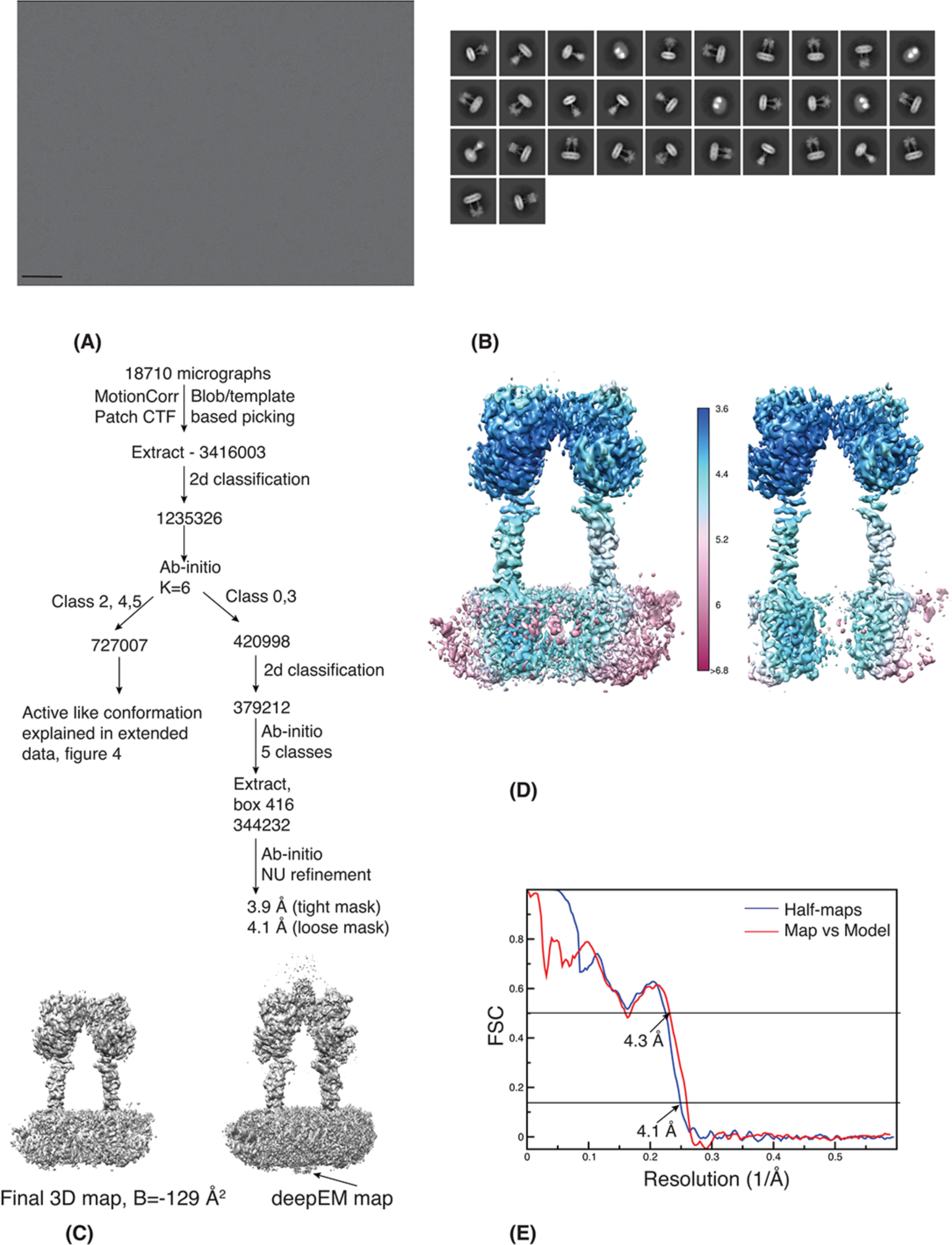
CryoEM analysis of quisqualate bound to Full-lenght mGlu_5_ (mGlu_5_-Δ856) in a Rcc conformation. (A) A representative micrograph of the quisqualate FL mGlu_5_ in detergent micelles imaged on ice and collected on K3 detector. Scale bar is 500 Å. (B) Selected reference-free 2D class averages of Rcc conformation of mGlu_5_ with a box size of 416 pixels and sampled at 0.84 Å. The separated TMD can be clearly seen in the class averages. (C) The image processing workflow of the mGlu_5_ with all the steps performed in cryosparc. The final 3D maps with either sharpened with a B-factor of −129 Å^2^ and generated with deepEMhancer using the half-maps and a mask are shown. The arrow marks the ICL2 loop in the deepEM maps that was used for model building. D) CryoEM maps coloured according to the local resolution plot of FL mGlu_5_ as determined with Relion 4.0 are shown in two different views. Much of the VFT in one monomer are at higher resolution and due to dynamic nature, the other monomer is of lower resolution. The CRD and the TMD in both monomers are at lower resolution along with the detergent-lipid micelle. (D) The Fourier shell correlation (FSC) curves of two half-maps with a mask (blue) and the map and model (red) are shown. The estimated resolution by comparison of the two half-maps from refinement and postprocessing at FSC 0.143 is 4.1 Å (the tight mask after corrections in cryosparc estimates 3.9 Å) and the one of map vs model at FSC 0.5 is 4.3 Å.

**Extended Data Figure 7.**
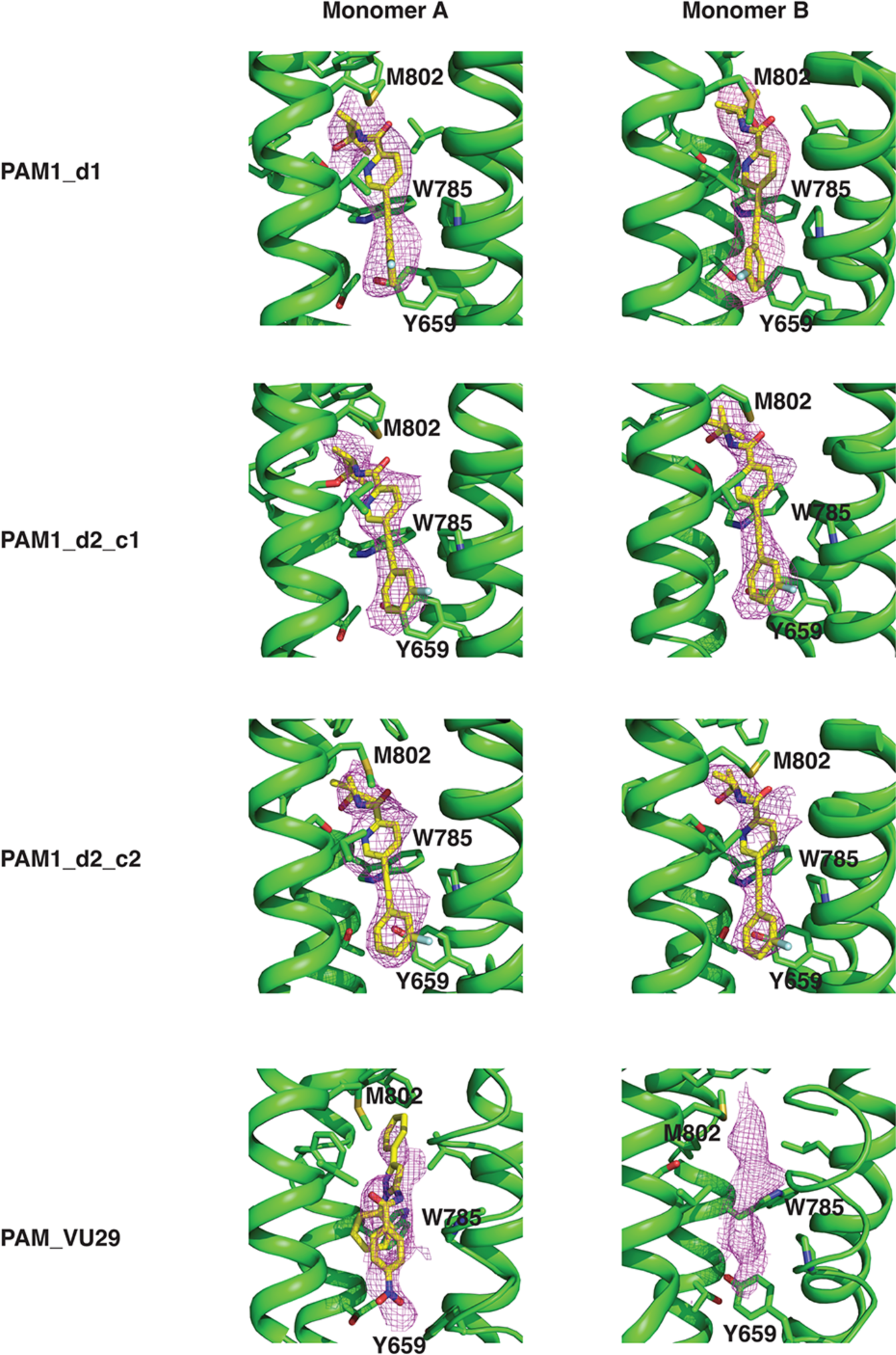
Omit maps of PAM in mGlu_5_ cryoEM maps. The unsharpened maps and the model without ligand from each data set was given as input in servalcat (Yamashita et al 2021) and the fofc_differencemap.mrc was used to demonstrate the presence of ligands as shown in the figure. In PAM, VU29, only one PAM (monomer A) has been modelled but to show the residual density in monomer B, the ligand was used as dummy to carve the density. The carbon atoms in protein are shown in green and cartoon representation and side chains of key residues are shown. The ligands are shown in stick representation and carbon atoms in yellow. The density (magenta) of the omit map was generated with Pymol carved around 2 Å with map_double option. The sigma values for the maps are: PAM1_d1 (7), PAM1_d2_c1 (5), PAM1_d2_c2 (7) and PAM_VU29 (3).

**Extended Data Figure 8.**
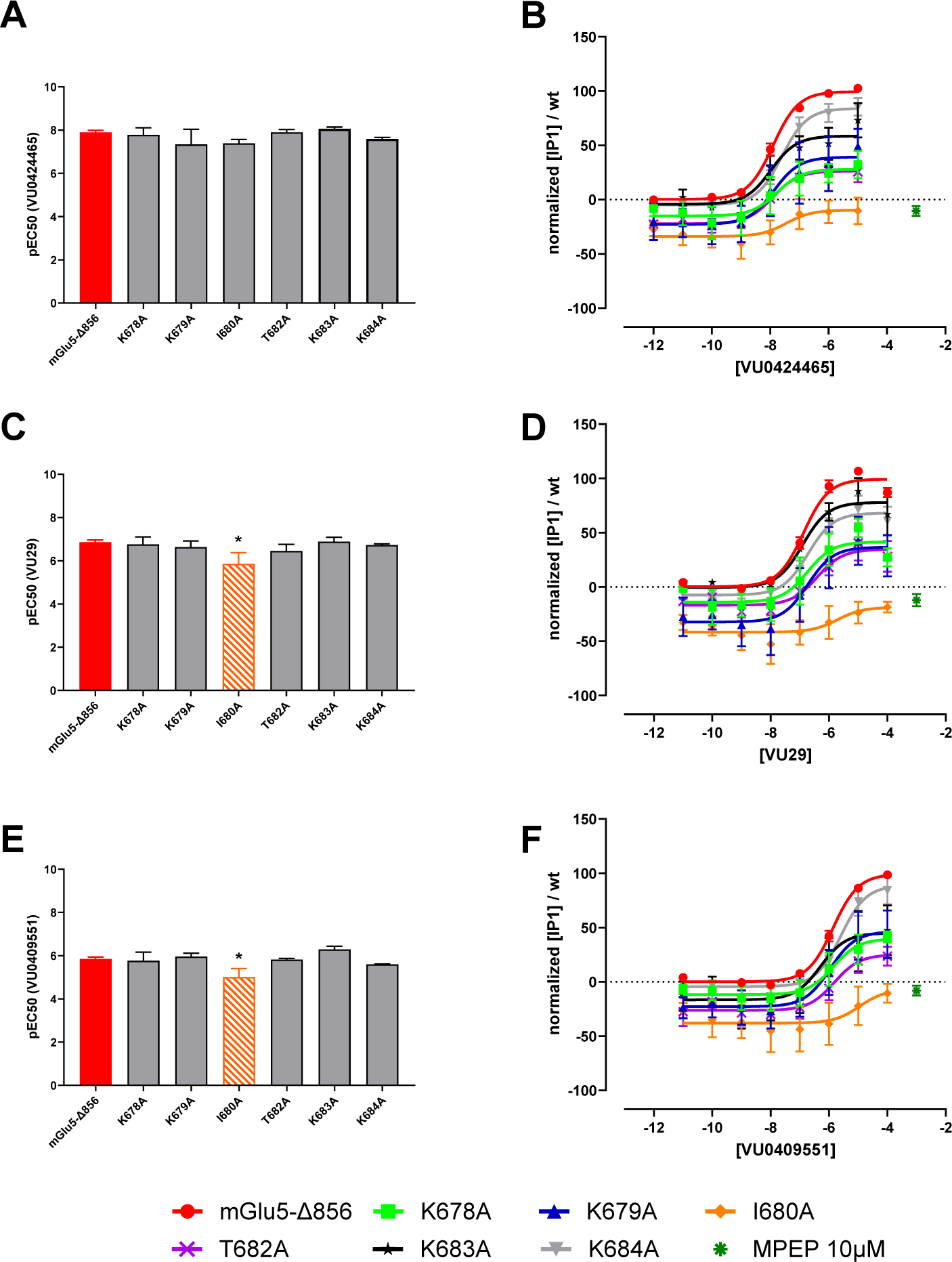
Functional consequences of mutations in the second intracellular loops (ICL2) of mGlu_5_ on PAM activity. Six different residues from the ICL2 of mGlu_5_-Δ856 were mutated into alanine by site-directed mutagenesis. The influence of these mutations on the dose-dependent potentiation of agonist-induced response was compared to the mGlu_5_-Δ856 receptor, using the IP-One HTRF assay in HEK293 cells transiently transfected with the indicated receptor. A, C, E. Comparison of the mean pEC50 ± SEM of VU0424465, VU29 and VU0409551, as determined from 3 independent dose-response experiments for all mutants except for K684A (n=4) and for mGlu_5_-Δ856 (n=7), in presence of a fixed concentration of the agonist quisqualate (10 nM) and different concentrations of the indicated PAM. B, D, F. Dose-dependent potentiation by VU0424465, VU29 and VU0409551 of the response induced by 10 nM of the agonist quisqualate on mGlu_5_-Δ856 and the mutants. Each data point corresponds to mean ± SEM of 3 independent experiments for all mutants except for K684A (n=4) and for mGlu_5_-Δ856 (n=7). Data are normalized to the response measured on the mGlu_5_-Δ856 receptor. Concentration-response curve parameters are listed in Supplementary Table 3. Difference between mGlu_5_-Δ856 and mutants were analyzed by a One-way ANOVA followed by Dunnett’s post hoc test. * p<0.05.

**Extended Data Figure 9.**
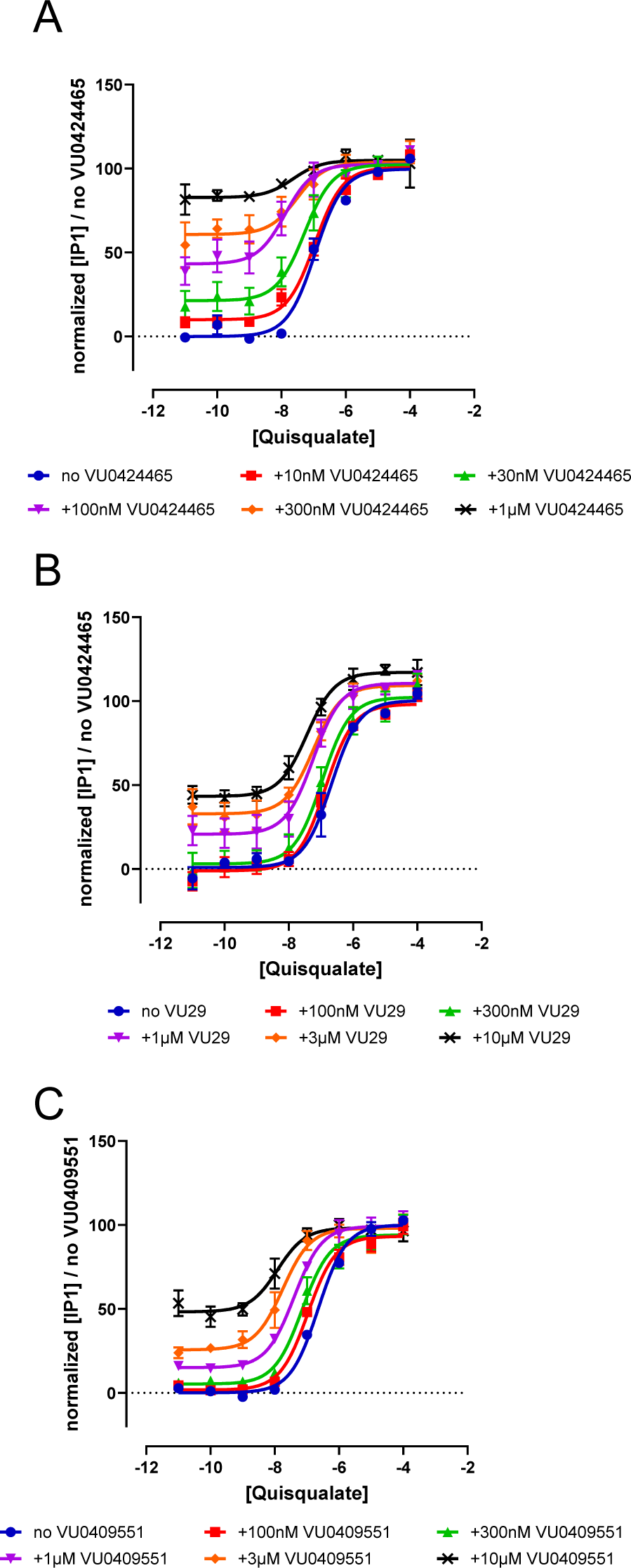
Dose-dependent potentiation of agonist-induced response of mGlu_5_ by VU0424465, VU29 and VU0409551. The ability of the PAMs to positively modulate the dose-dependent response of mGlu_5_ to the agonist quisqualate was determined by the measurement of IP_1_ accumulation in HEK cells transiently transfected with mGlu_5_-Δ856. A. Effect of VU0424465. B. Effect of VU29. C. Effect of VU0409551. Data are normalized to the response measured for quisqualate alone and are mean ± SEM of at least three independent experiments performed in duplicate. Parameters from the dose-response curves are listed in Supplementary Table 4.

**Extended Data Figure 10.**
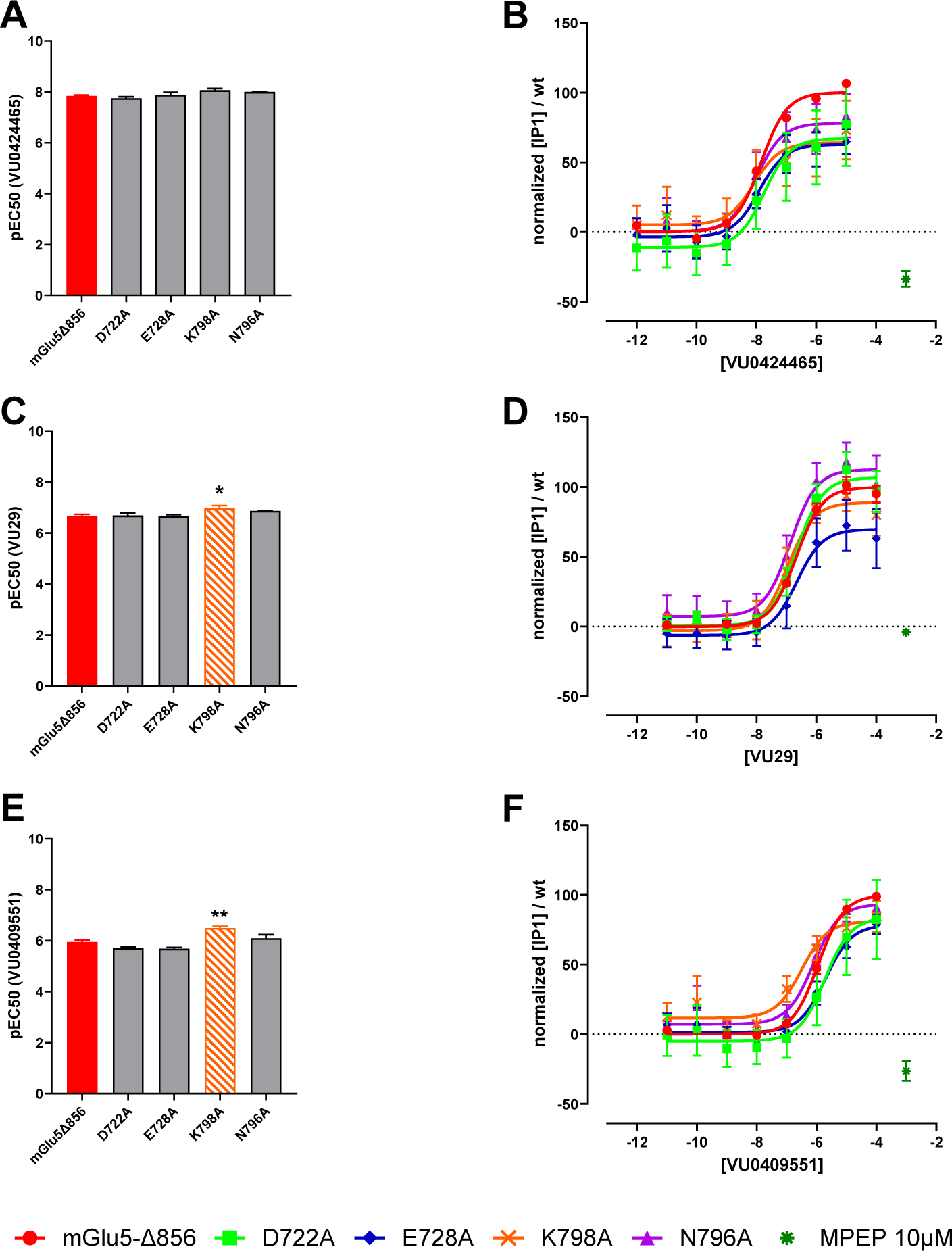
Functional consequences of mutations in the second and third extracellular loop (ECL2, ECL3) of mGlu_5_ on PAM activity. Three different residues from the ECL2 and one residue of the ECL3 of mGlu_5_ were mutated into alanine by site-directed mutagenesis. The influence of these mutations on the dose-dependent potentiation of agonist-induced response was compared to the mGlu_5_-Δ856 receptor, using the IP-One HTRF assay. Comparison of the mean pEC50 ± SEM of VU0424465, VU29 and VU0409551, as determined from three independent dose-response experiments in presence of a fixed concentration of the agonist Quisqualate (10 nM) and different concentrations of the indicated PAM (A, C and E). Dose-dependent potentiation by VU0424465, VU29 and VU0409551 of the response induced by 10 nM of the agonist quisqualate of mGlu_5_-Δ856 and mutants (B, D and F). Each data point corresponds to mean ± SEM of three independent experiments. Data are normalized to the response measured on the WT receptor. Concentration-response curve parameters are listed in Supplementary Table 2. Difference between mGlu_5_-Δ856 and mutants were analyzed by a One-way ANOVA followed by Dunnett’s post hoc test. * p<0.05, ** p<0.01.

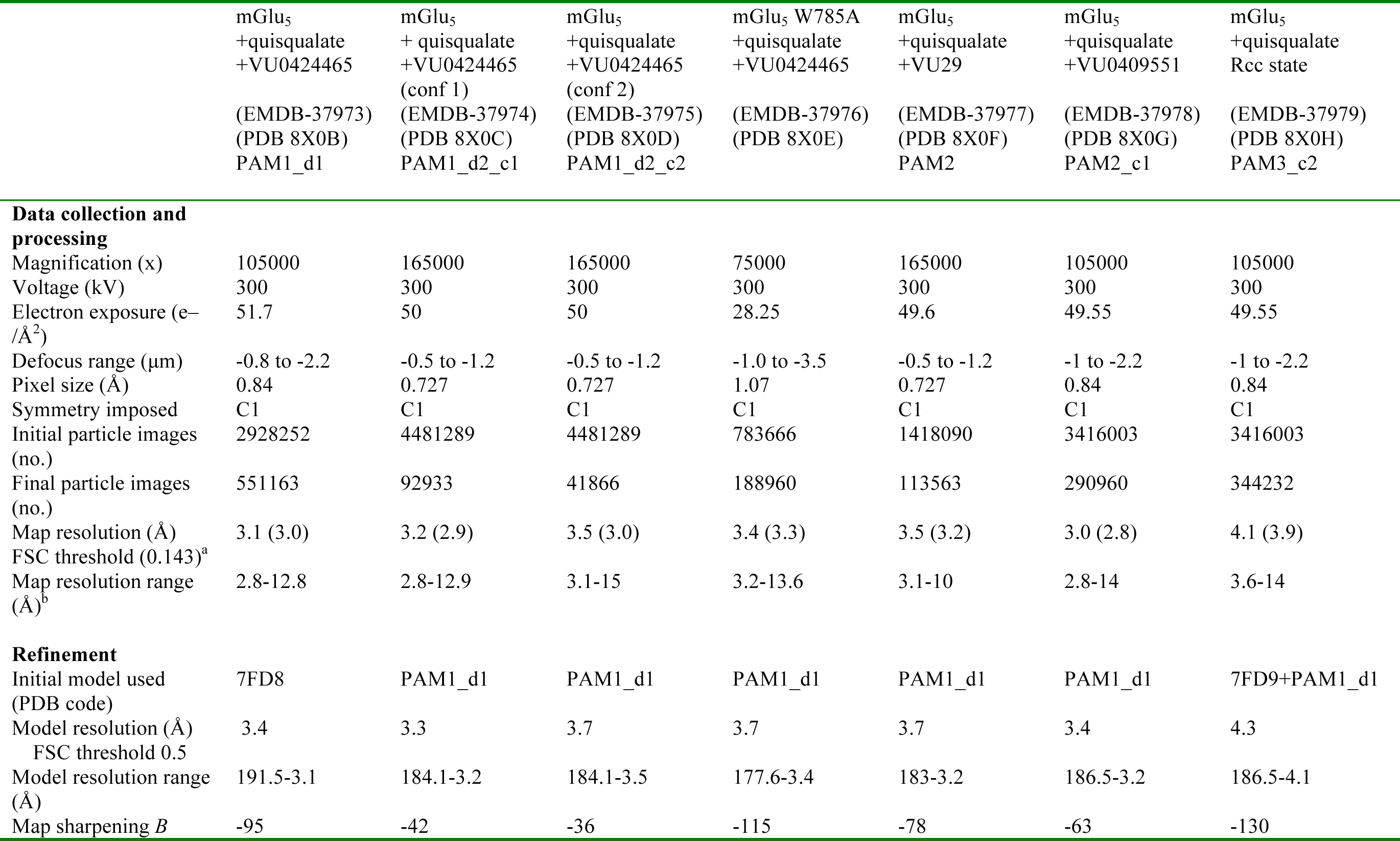
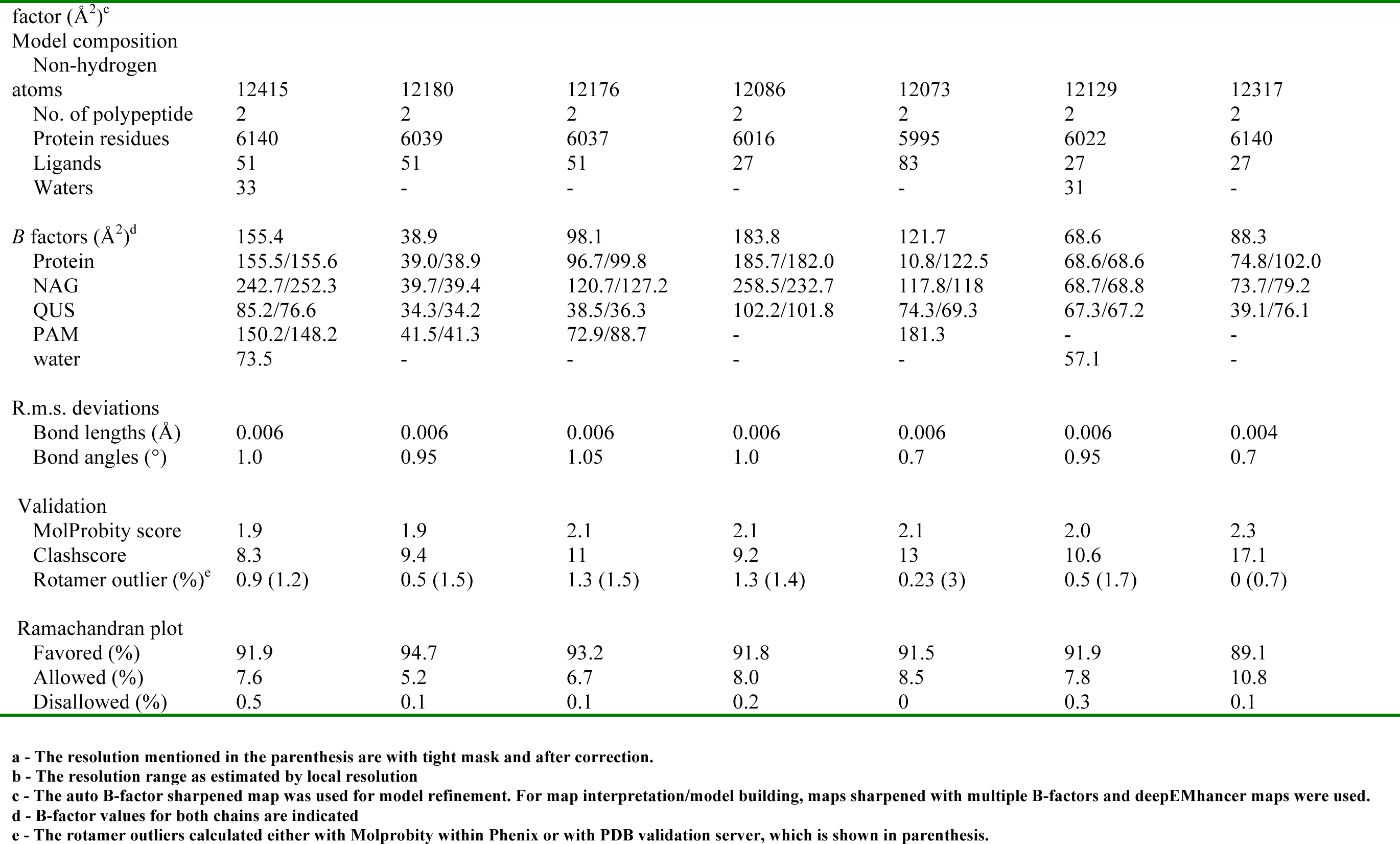
Cryo-EM data collection, refinement and validation statistics.

**Supplementary Figure 1.**
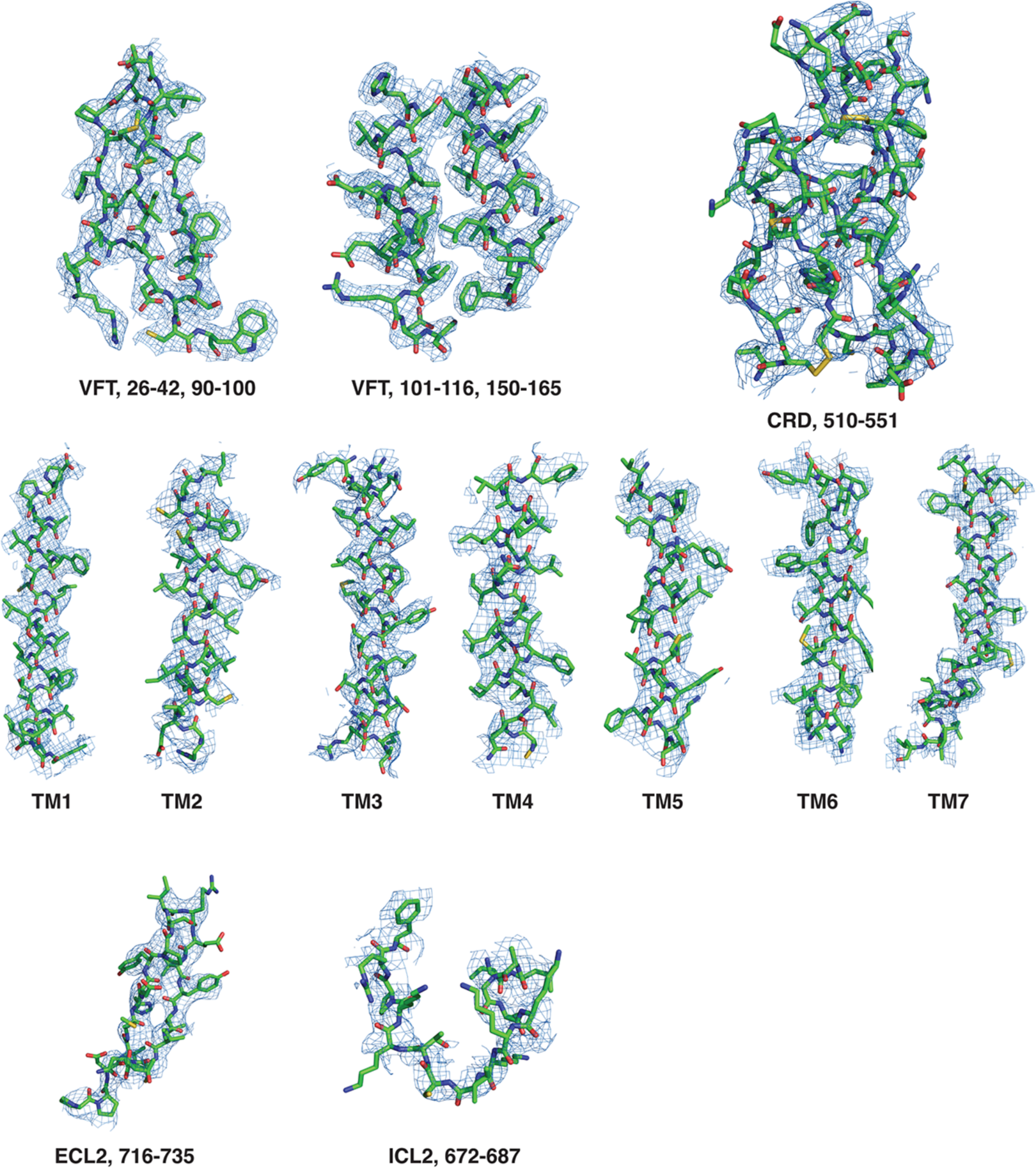
Example density of select regions of the cryoEM map of PAM1_d1 of mGlu_5_. The cryoEM map is shown in blue mesh and the receptor in stick representation. The regions of the VFT and the CRD are shown in the top panel using the auto B-factor sharpened map at 8 σ. The CRD density was made with map sharpened with B factor of −45 Å^2^ and at 10.5σ In the middle panel, the densities for the 7TM are shown using the map sharpened with B factor of −45 Å^2^ (TM 1,2, 4,6,7 at 8σ, TM3 at 8.5 σ and TM5 at 9 σ). The ECL2 density (11 σ) was also made with map sharpened with B factor of −45 Å^2^. For the ICL2 region, deepEMhancer map was used in model building and shown here.

**Supplementary Figure 2.**
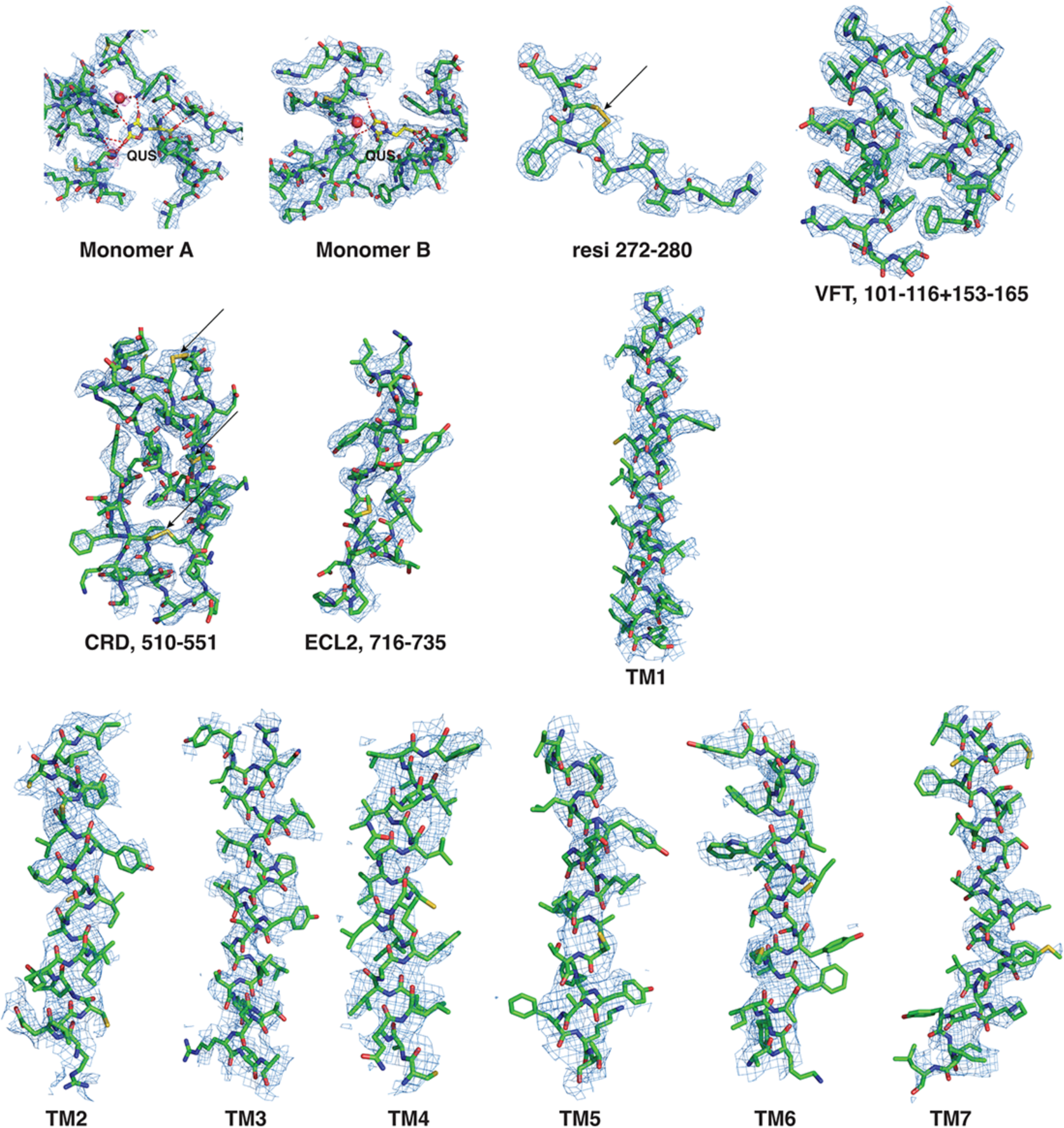
Example density of select regions of the cryoEM maps of PAM3_c1 of mGlu_5_. The cryoEM map is shown in blue mesh, the receptor in stick representation and the water molecules in red sphere. In the top panel, regions around the quisqualate binding sites in both monomers and the VFT are shown. The disulphide is marked with an arrow. The sharpened map with automatic B-factor was used to make this map at 8 σ. The density for water is shown in magenta. In the middle panel, the CRD with disulphides marked with arrow, ECL2 (8σ) and TM1 are shown. Of the 10 disulphide bonds in each monomer, good density is observed for 7 of them i.e., observable beyond radiation damage. In the bottom panel, density for TM helices 2-7 are shown. The map sharpened with B=-30 Å^2^ was used to make the maps for TM helices (TM 1 – 5.5 σ, TM 2,3 – 6 σ, TM 4,5 – 6.5 σ, TM 6 – 7 σ and TM 7 – 6.5 σ).

**Supplementary Figure 3.**
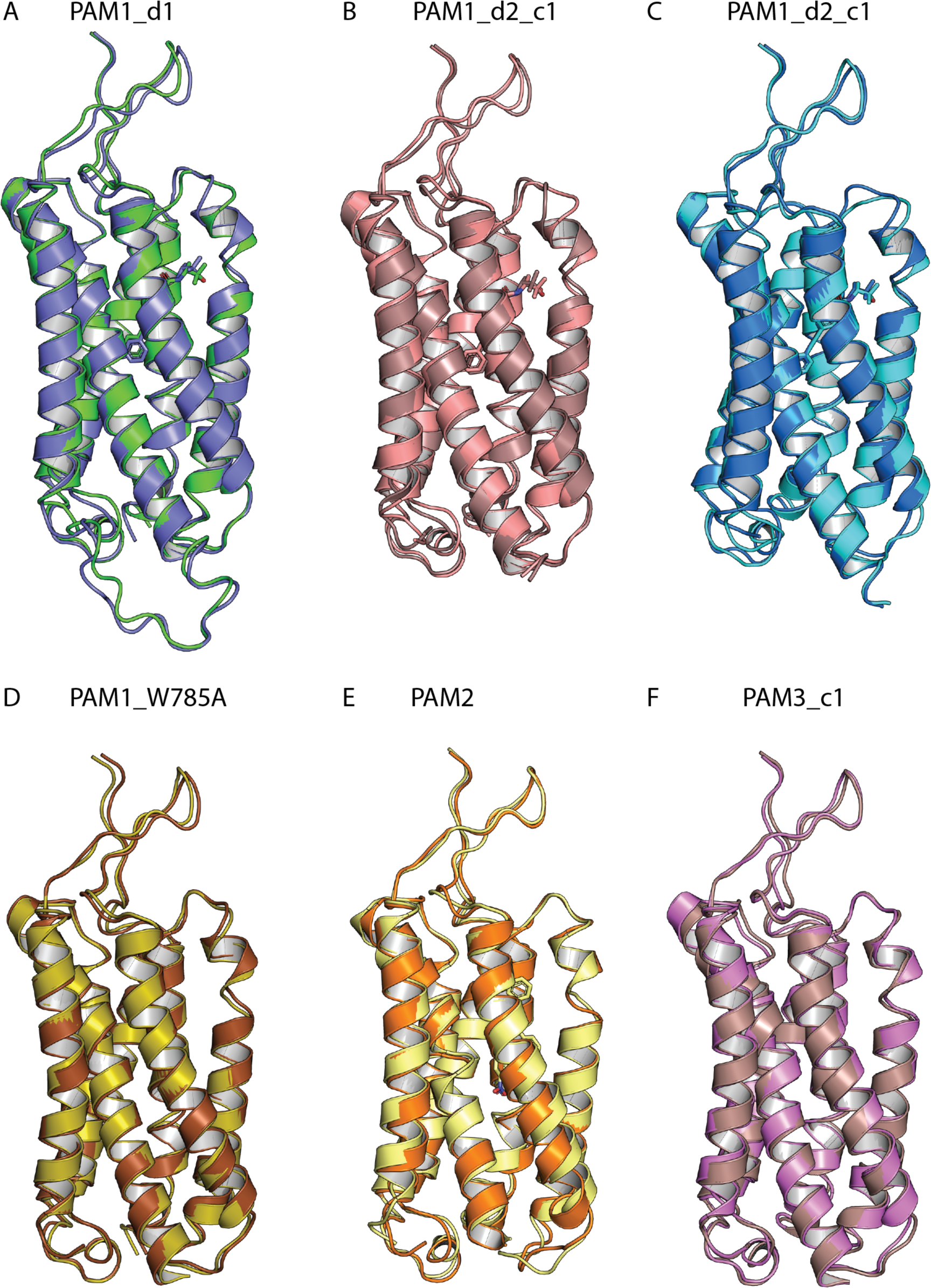
Superimposition of the two 7TM domains from each determined Acc structure. VU0424465 models (PAM1 data sets) display a shifted of H6 towards the extracellular side for one protomer and a kink at P790. Shift of TM6 and the kink at P790 is much less pronounced for W785A mutant and for the two other structures (PAM2 and PAM3_d1).

**Supplementary Figure 4.**
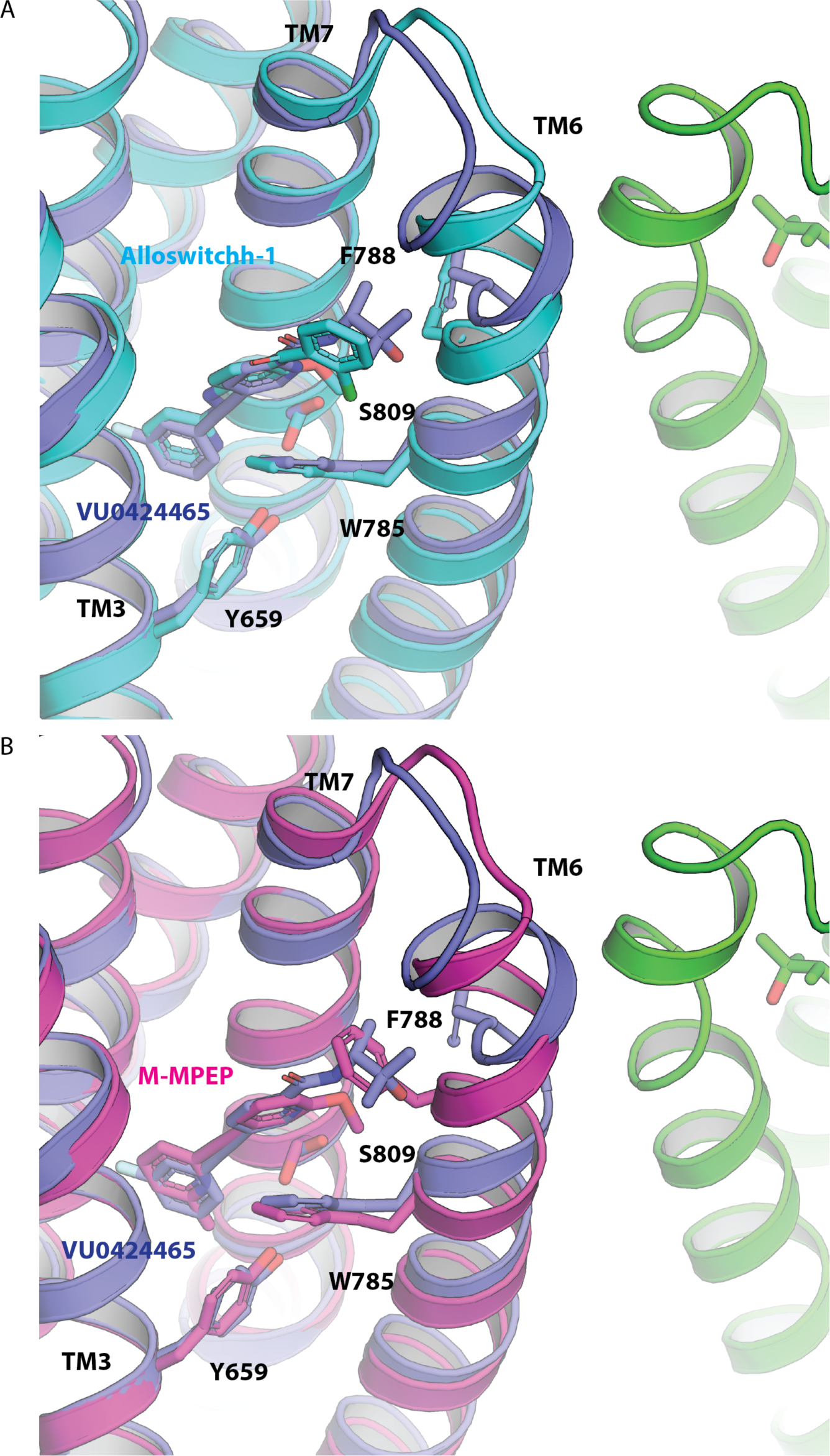
PAMs and NAMs occupy a common binding site. Superposition of the X-ray structure of thermostabilized mGlu_5_ 7TM (mGlu_5_-StaR(569-836)-T4L) bound to NAM alloswitch-1 (A; Cyan, PDB code 7P2L) and M-MPEP (B; Purple, PDB code 6FFI) with 7TM of the mGlu_5_-5M bound to ago-PAM VU0424465 (blue and green) determined in this work.

**Supplementary Figure 5.**
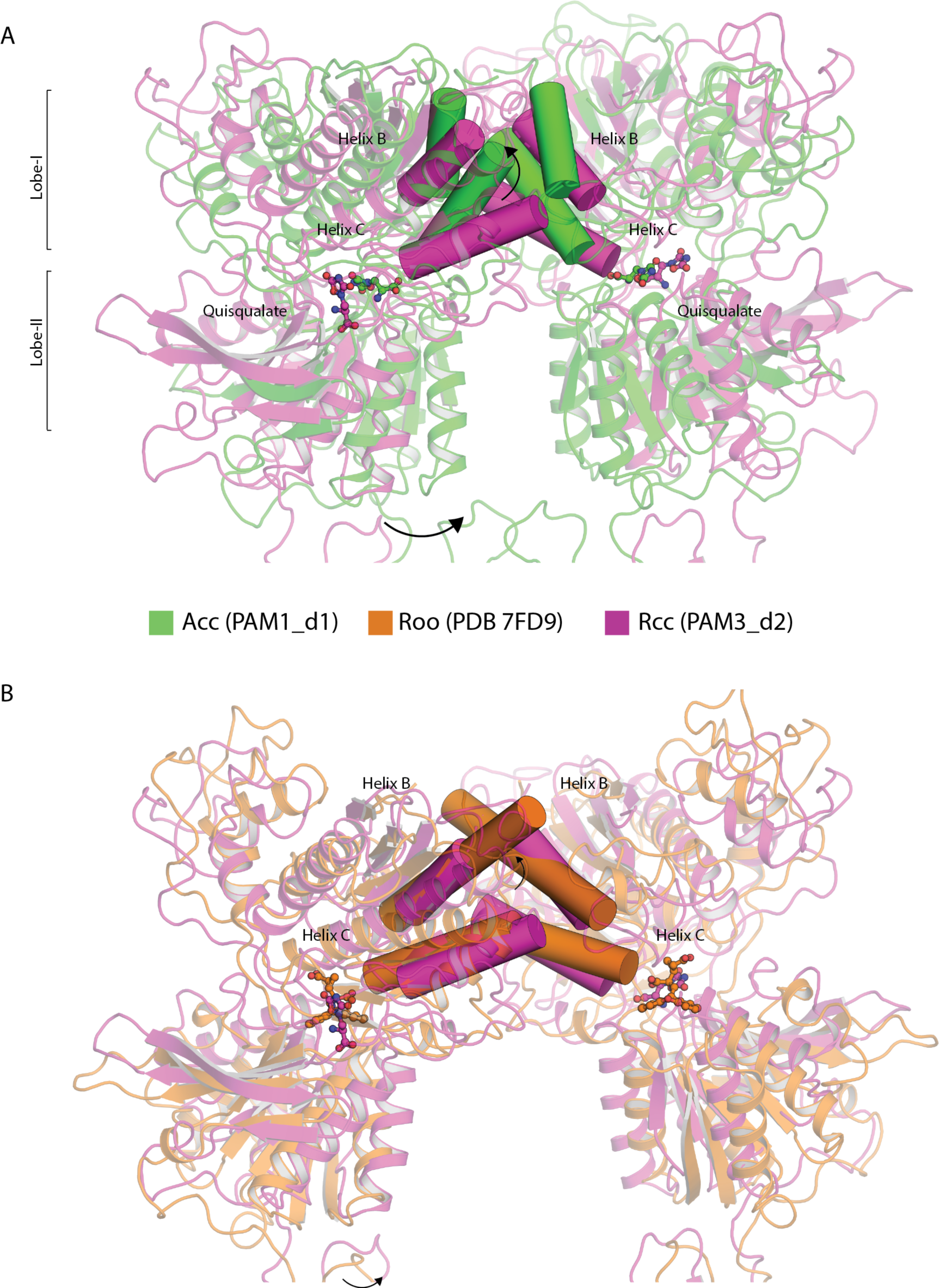
Comparison of the Rcc Intermediate-active state with Acc fully active and Roo fully inactive state of the VFTs. Rcc displays closed VFTs in resting R state similar to the fully inactive state Roo. Although VFTs are in close conformation, lobe-I did not rotate (A). Rcc conformation is structurally similar to the Roo fully inactive state of the VFTs, however, Helix B and C that mediates the Lobes-1 interactions started to shift toward the active state.

**Supplementary Figure 6.**
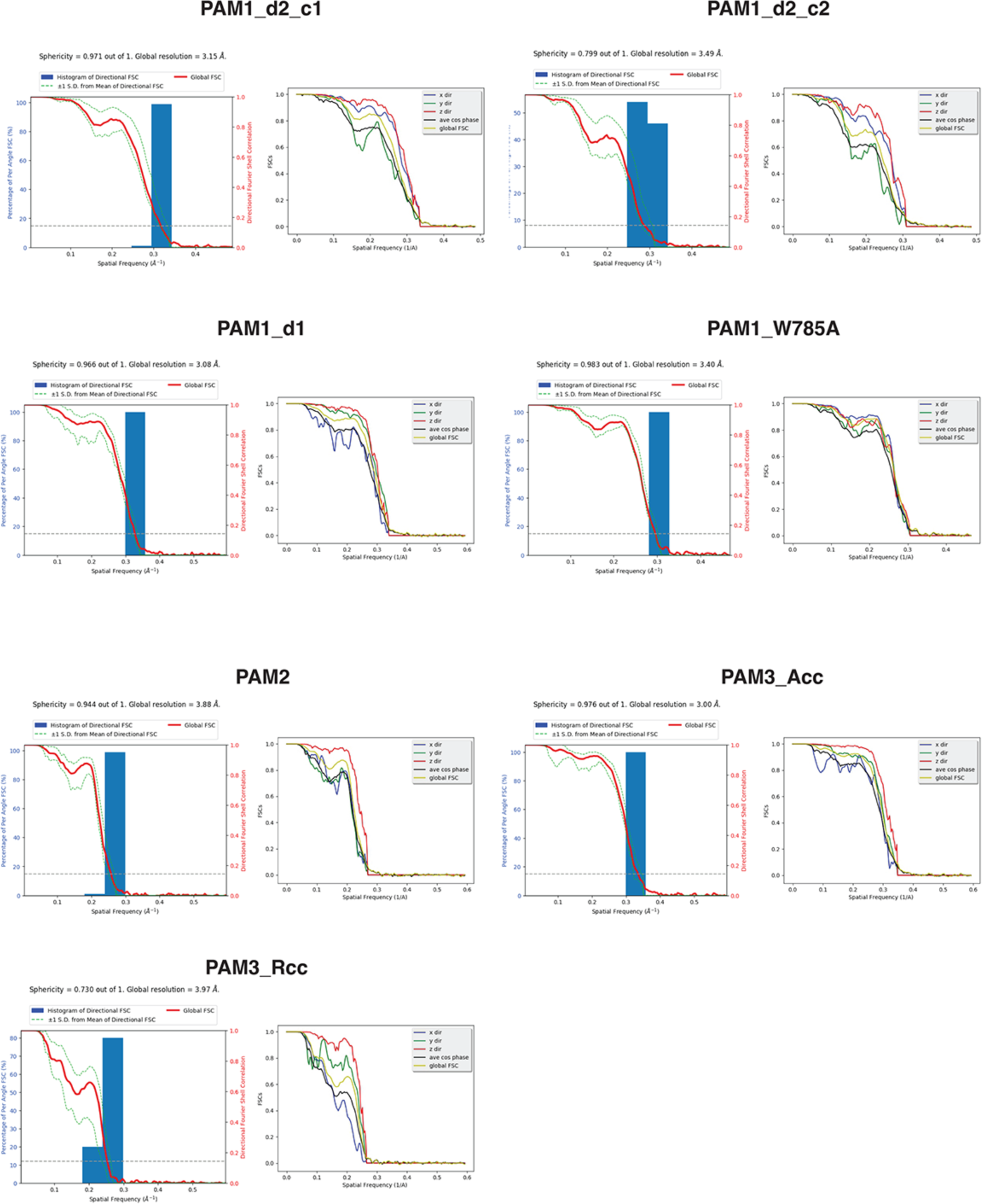
3D FSC analysis of all the data sets of mGlu_5_ from this study. The histogram of directional FSC’s as calculated by the 3D FSC server (Tan et al 2017) for the 7 data sets are shown. Except for the Pam1_d2_c2 and PAM3_Rcc, all the data show high sphericity.

**Supplementary Table 1.** Comparison of the potencies of the dose-dependent potentiation of agonist-induced response of VU0424465, VU29 and VU0409551 on mGlu_5_-Δ856 and different mutants in the intracellular loop 2. Potencies were determined by fitting individual dose-dependent curves of the potentiation by VU0424465, VU29 or VU0409551 of the production of inositol monophosphate induced by a fixed concentration of the agonist quisqualate (10 nM) on HEK293 cells transiently expressing either mGlu_5_-Δ856 or mGlu_5_-Δ856 carrying different single alanine mutations in the second intracellular loop (ICL2). The data correspond to the values of the histograms shown in Extended Data Figure 8 and are mean ± SEM of ‘n’ independent experiments. Difference between potencies measured on mGlu5-Δ856 and various mutants were analyzed by a One-way ANOVA followed by Dunnett’s post hoc test. * p<0.05.

|  | **VU0424465**<br>**pEC50±SEM (n)** | **VU29**<br>**pEC50±SEM (n)** | **VU0409551**<br>**pEC50±SEM (n)** |
| --- | --- | --- | --- |
| mGlu<sub>5</sub>-Δ856 | 7,9±0,10 (7) | 6,86±0,10 (7) | 5,85±0,09 (7) |
| K678A | 7,78±0,33 (3) | 6,76±0,34 (3) | 5,77±0,39 (3) |
| K679A | 7,34±0,70 (3) | 6,63±0,28 (3) | 5,96±0,15 (3) |
| I680A | 7,39±0,18 (3) | 5,86±0,52 (3) * | 5,01±0,39 (3) * |
| T682A | 7,90±0,13 (3) | 6,46±0,31 (3) | 5,82±0,06 (3) |
| K683A | 8,06±0,09 (3) | 6,89±0,20 (3) | 6,29±0,15 (3) |
| K684A | 7,59±0,07 (4) | 6,73±0,05 (4) | 5,60±0,02 (4) |

**Supplementary Table 2.** Comparison of the potencies, basal and maximal efficacies of the agonist quisqualate in presence of fixed concentrations of VU0424465, VU29 and VU0409551 on mGlu_5_-Δ856. Potencies, basal (Ebasal) and maximal efficacies (Emax) were determined by fitting the dose-dependent production of inositol monophosphate induced by quisqualate, in absence and in presence of fixed concentrations of VU0424465, VU29 or VU0409551 on HEK293 cells transiently expressing mGlu_5_-Δ856. Efficacy values is normalized to the basal and maximal efficacy of quisqualate in absence of PAM. The data correspond to the curves shown in Extended Data Figure 9 and are mean ± SEM of three independent experiments.

|  | pEC50 |  | E basal |  | E max |  | n |
| --- | --- | --- | --- | --- | --- | --- | --- |
|  | mean | SEM | mean | SEM | mean | SEM |  |
| [VU0424465] |  |  |  |  |  |  |  |
| no VU | -6,93 | 0,09 | 0,00 | 0,00 | 100,00 | 0,00 | 3 |
| 10nM | -6,94 | 0,08 | 10,01 | 2,23 | 100,96 | 3,49 | 3 |
| 30nM | -7,29 | 0,21 | 20,77 | 7,65 | 102,65 | 4,44 | 3 |
| 100nM | -7,84 | 0,23 | 43,55 | 7,83 | 103,39 | 2,55 | 3 |
| 300nM | -7,50 | 0,35 | 60,80 | 8,20 | 105,03 | 0,61 | 3 |
| 1µM | -6,84 | 1,29 | 83,11 | 6,09 | 114,74 | 12,88 | 3 |
| [VU29] |  |  |  |  |  |  |  |
| no VU | -6,72 | 0,17 | 0,00 | 0,00 | 100,00 | 0,00 | 3 |
| 100nM | -6,85 | 0,05 | -1,14 | 5,11 | 98,23 | 2,68 | 3 |
| 300nM | -6,96 | 0,05 | 3,00 | 7,38 | 102,61 | 7,89 | 3 |
| 1µM | -7,24 | 0,11 | 20,48 | 9,16 | 110,73 | 3,69 | 3 |
| 3µM | -7,25 | 0,05 | 32,70 | 8,45 | 109,23 | 4,47 | 3 |
| 10µM | -7,43 | 0,17 | 42,95 | 4,15 | 118,03 | 1,08 | 3 |
| [VU0409551] |  |  |  |  |  |  |  |
| no VU | -6,65 | 0,02 | 0,00 | 0,00 | 100,00 | 0,00 | 3 |
| 100nM | -6,98 | 0,07 | 1,63 | 1,06 | 93,22 | 5,02 | 3 |
| 300nM | -7,13 | 0,12 | 5,19 | 1,45 | 94,35 | 6,44 | 3 |
| 1µM | -7,40 | 0,04 | 14,94 | 0,81 | 99,37 | 6,34 | 3 |
| 3µM | -7,78 | 0,21 | 24,72 | 1,09 | 98,12 | 1,51 | 3 |
| 10µM | -7,99 | 0,22 | 46,23 | 4,38 | 98,77 | 3,51 | 3 |

**Supplementary Table 3.** Comparison of the potencies of the dose-dependent potentiation of agonist-induced response of VU0424465, VU29 and VU0409551 at mGlu_5_-Δ856 and selected mutants in PAM the binding site. Potencies were determined by fitting individual dose-dependent curves of the potentiation by VU0424465, VU29 or VU0409551 of the production of inositol monophosphate induced by a fixed concentration of the agonist quisqualate (10 nM) on HEK293 cells transiently expressing either mGlu_5_-Δ856 or mGlu_5_-Δ856 carrying different single alanine mutations in the 7TM domain. The data correspond to the values of the histograms shown in Figure 4 and are mean ± SEM of n-independent experiments. Difference between potencies measured on mGlu_5_-Δ856 and various mutants were analyzed by a One-way ANOVA followed by Dunnett’s post hoc test. * p<0.05, ** p<0.01, *** p<0.005, **** p<0.001.

|  | VU0424465<br>pEC50±SEM (n) | VU29<br>pEC50±SEM (n) | VU0409551<br>pEC50±SEM (n) |
| --- | --- | --- | --- |
| mGlu<sub>5</sub>-Δ856 | 7,93±0,07 (8) | 6,87±0,11 (7) | 5,95±0,11 (8) |
| S658A<sup>3.43</sup> | 8,27±0,03 (4) | 7,19±0,01 (3) | 6,4±0,2 (4) |
| Y659A<sup>3.44</sup> | 5,68±0,02 (3) **** | 2±1,92 (3) **** | 4,93±0,17 (4) *** |
| T781A<sup>6.86</sup> | 6,56±0,09 (4) **** | 6,55±0,08 (4) | 5,0±0,28 (4) ** |
| W785A<sup>6.50</sup> | 7,98±0,11 (4) | 8,33±0,1 (4) **** | 5,14±0,12 (4) ** |
| F788A<sup>6.53</sup> | 8,34±0,02 (3) | 5,50±0,13 (3) **** | 6,58±0,12 (3) |
| Y792A<sup>6.57</sup> | 7,68±0,37 (4) | 5,97±0,08 (3) **** | 6,24±0,26 (4) |
| F793A<sup>6.58</sup> | 7,90±0,34 (4) | 6,73±0,06 (3) | 6,33±0,24 (4) |
| S805A<sup>7.35</sup> | 7,86±0,07 (4) | 6,77±0,06 (4) | 5,96±0,06 (4) |
| S809A<sup>7.39</sup> | 6,92±0,08 (4) *** | 5,42±0,14 (4) **** | 5,03±0,19 (4) |

**Supplementary Table 4.** Comparison of the potencies of the dose-dependent potentiation of agonist-induced response of VU0424465, VU29 and VU0409551 on mGlu_5_-Δ856 and different mutants of the second and third extracellular loop. Potencies were determined by fitting individual dose-dependent curves of the potentiation by VU0424465, VU29 or VU0409551 of the production of inositol monophosphate induced by a fixed concentration of the agonist quisqualate (10 nM) on HEK293 cells transiently expressing either mGlu_5_-Δ856 or mGlu_5_-Δ856 carrying different single alanine mutations in the second and third extracellular loop (ECL2, ECL3). The data correspond to the values of the histograms shown in Extended Data Figure 10 and are mean ± SEM of n-independent experiments. Difference between potencies measured on mGlu_5_-Δ856 and various mutants were analyzed by a One-way ANOVA followed by Dunnett’s post hoc test. * p<0.05, ** p<0.01.

|  | **VU0424465**<br>pEC50±SEM (n) | **VU29**<br>pEC50±SEM (n) | **VU0409551**<br>pEC50±SEM (n) |
| --- | --- | --- | --- |
| mGlu<sub>5</sub>-Δ856 | 7,84±0,03 (3) | 6,66±0,07 (3) | 5,95±0,08 (3) |
| D722A<br>(ECL3) | 7,75±0,06 (3) | 6,69±0,10 (3) | 5,71±0,05 (3) |
| E728A<br>(ECL3) | 7,88±0,10 (3) | 6,66±0,06 (3) | 5,69±0,05 (3) |
| N796A(ECL2) | 8,00±0,02 (3) | 6,87±0,00 (3) * | 6,10±0,14 (3) ** |
| K798A(ECL2) | 8,06±0,07 (3) | 6,98±0,10 (3) | 6,50±0,07 (3) |

## References

1. Sladeczek, F., Pin, J. P., Récasens, M., Bockaert, J. & Weiss, S. Glutamate stimulates inositol phosphate formation in striatal neurones. Nature 317, 717–719 (1985).

2. Pin, J.-P. & Bettler, B. Organization and functions of mGlu and GABAB receptor complexes. Nature 540, 60–68 (2016).

3. Wang, X. et al. Structural insights into dimerization and activation of the mGlu2-mGlu3 and mGlu2-mGlu4 heterodimers. Cell Res 33, 762–774 (2023).

4. Seven, A. B. et al. G-protein activation by a metabotropic glutamate receptor. Nature 595, 450–454 (2021).

5. Lin, S. et al. Structures of Gi-bound metabotropic glutamate receptors mGlu2 and mGlu4. Nature 594, 583–588 (2021).

6. Koehl, A. et al. Structural insights into the activation of metabotropic glutamate receptors. Nature 566, 79–84 (2019).

7. Gregory, K. J. & Goudet, C. International Union of Basic and Clinical Pharmacology. CXI. Pharmacology, Signaling, and Physiology of Metabotropic Glutamate Receptors. Pharmacol Rev 73, 521–569 (2021).

8. Stansley, B. J. & Conn, P. J. Neuropharmacological Insight from Allosteric Modulation of mGlu Receptors. Trends Pharmacol Sci 40, 240–252 (2019).

9. Dogra, S. & Conn, P. J. Metabotropic Glutamate Receptors As Emerging Targets for the Treatment of Schizophrenia. Mol Pharmacol 101, 275–285 (2022).

10. Du, J. et al. Structures of human mGlu2 and mGlu7 homo- and heterodimers. Nature 594, 589–593 (2021).

11. Nasrallah, C. et al. Agonists and allosteric modulators promote signaling from different metabotropic glutamate receptor 5 conformations. Cell Rep 36, 109648 (2021).

12. Nicoletti, F. et al. Metabotropic glutamate receptors: from the workbench to the bedside. Neuropharmacology 60, 1017–1041 (2011).

13. Parmentier-Batteur, S. et al. Mechanism based neurotoxicity of mGlu5 positive allosteric modulators--development challenges for a promising novel antipsychotic target. Neuropharmacology 82, 161–173 (2014).

14. Rook, J. M. et al. Unique signaling profiles of positive allosteric modulators of metabotropic glutamate receptor subtype 5 determine differences in in vivo activity. Biol Psychiatry 73, 501–509 (2013).

15. Rook, J. M. et al. Biased mGlu5-Positive Allosteric Modulators Provide In Vivo Efficacy without Potentiating mGlu5 Modulation of NMDAR Currents. Neuron 86, 1029– 1040 (2015).

16. Sengmany, K. et al. Biased allosteric agonism and modulation of metabotropic glutamate receptor 5: Implications for optimizing preclinical neuroscience drug discovery. Neuropharmacology 115, 60–72 (2017).

17. Hellyer, S. D. et al. Probe dependence and biased potentiation of metabotropic glutamate receptor 5 is mediated by differential ligand interactions in the common allosteric binding site. Biochem Pharmacol 177, 114013 (2020).

18. Chen, Y. et al. Interaction of novel positive allosteric modulators of metabotropic glutamate receptor 5 with the negative allosteric antagonist site is required for potentiation of receptor responses. Mol Pharmacol 71, 1389–1398 (2007).

19. Sengmany, K. et al. Differential contribution of metabotropic glutamate receptor 5 common allosteric binding site residues to biased allosteric agonism. Biochem Pharmacol 177, 114011 (2020).

20. Gregory, K. J. et al. Investigating Metabotropic Glutamate Receptor 5 Allosteric Modulator Cooperativity, Affinity, and Agonism: Enriching Structure-Function Studies and Structure-Activity Relationships. Mol Pharmacol 82, 860–875 (2012).

21. Noetzel, M. J. et al. Functional impact of allosteric agonist activity of selective positive allosteric modulators of metabotropic glutamate receptor subtype 5 in regulating central nervous system function. Mol Pharmacol 81, 120–133 (2012).

22. Park, J. et al. Symmetric activation and modulation of the human calcium-sensing receptor. Proceedings of the National Academy of Sciences 118, e2115849118 (2021).

23. Shen, C. et al. Structural basis of GABAB receptor–Gi protein coupling. Nature 594, 594–598 (2021).

24. Gao, Y. et al. Asymmetric activation of the calcium-sensing receptor homodimer. Nature 595, 455–459 (2021).

25. Lecat-Guillet, N. et al. Concerted conformational changes control metabotropic glutamate receptor activity. Sci Adv 9, eadf1378.

26. Doré, A. S. et al. Structure of class C GPCR metabotropic glutamate receptor 5 transmembrane domain. Nature 511, 557–562 (2014).

27. Christopher, J. A. et al. Fragment and Structure-Based Drug Discovery for a Class C GPCR: Discovery of the mGlu5 Negative Allosteric Modulator HTL14242 (3-Chloro-5-[6-(5-fluoropyridin-2-yl)pyrimidin-4-yl]benzonitrile). J Med Chem 58, 6653–6664 (2015).

28. Christopher, J. A. et al. Structure-Based Optimization Strategies for G Protein-Coupled Receptor (GPCR) Allosteric Modulators: A Case Study from Analyses of New Metabotropic Glutamate Receptor 5 (mGlu5) X-ray Structures. J. Med. Chem. 62, 207–222 (2019).

29. Nasrallah, C. et al. Direct coupling of detergent purified human mGlu5 receptor to the heterotrimeric G proteins Gq and Gs. Sci Rep 8, 4407 (2018).

30. Sanchez-Garcia, R. et al. DeepEMhancer: a deep learning solution for cryo-EM volume post-processing. Commun Biol 4, 874 (2021).

31. Gregory, K. J. et al. Identification of Specific Ligand–Receptor Interactions That Govern Binding and Cooperativity of Diverse Modulators to a Common Metabotropic Glutamate Receptor 5 Allosteric Site. ACS Chem Neurosci 5, 282–295 (2014).

32. Vafabakhsh, R., Levitz, J. & Isacoff, E. Y. Conformational dynamics of a class C G protein-coupled receptor. Nature 524, 497–501 (2015).

33. Afonine, P. V. et al. Conformational space exploration of cryo-EM structures by variability refinement. Biochim Biophys Acta Biomembr 1865, 184133 (2023).

34. Gomeza, J. et al. The second intracellular loop of metabotropic glutamate receptor 1 cooperates with the other intracellular domains to control coupling to G-proteins. J Biol Chem 271, 2199–2205 (1996).

35. Goudet, C. et al. Heptahelical domain of metabotropic glutamate receptor 5 behaves like rhodopsin-like receptors. Proc Natl Acad Sci U S A 101, 378–383 (2004).

36. Gregory, K. J. et al. Probing the metabotropic glutamate receptor 5 (mGlu₅) positive allosteric modulator (PAM) binding pocket: discovery of point mutations that engender a ‘molecular switch’ in PAM pharmacology. Mol Pharmacol 83, 991–1006 (2013).

37. Arsova, A. et al. Positive Allosteric Modulators of Metabotropic Glutamate Receptor 5 as Tool Compounds to Study Signaling Bias. Mol Pharmacol 99, 328–341 (2021).

38. Venkatakrishnan, A. J. et al. Molecular signatures of G-protein-coupled receptors. Nature 494, 185–194 (2013).

39. Reeves, P. J., Callewaert, N., Contreras, R. & Khorana, H. G. Structure and function in rhodopsin: high-level expression of rhodopsin with restricted and homogeneous N-glycosylation by a tetracycline-inducible N-acetylglucosaminyltransferase I-negative HEK293S stable mammalian cell line. Proc Natl Acad Sci U S A 99, 13419–13424 (2002).

40. Punjani, A., Rubinstein, J. L., Fleet, D. J. & Brubaker, M. A. cryoSPARC: algorithms for rapid unsupervised cryo-EM structure determination. Nat Methods 14, 290–296 (2017).

41. Punjani, A., Zhang, H. & Fleet, D. J. Non-uniform refinement: adaptive regularization improves single-particle cryo-EM reconstruction. Nat Methods 17, 1214–1221 (2020).

42. Scheres, S. H. W. RELION: Implementation of a Bayesian approach to cryo-EM structure determination. J Struct Biol 180, 519–530 (2012).

43. Rohou, A. & Grigorieff, N. CTFFIND4: Fast and accurate defocus estimation from electron micrographs. J Struct Biol 192, 216–221 (2015).

44. Bepler, T. et al. Positive-unlabeled convolutional neural networks for particle picking in cryo-electron micrographs. Nat Methods 16, 1153–1160 (2019).

45. Zivanov, J. et al. New tools for automated high-resolution cryo-EM structure determination in RELION-3. Elife 7, e42166 (2018).

46. Weis, F. & Hagen, W. J. H. Combining high throughput and high quality for cryo-electron microscopy data collection. Acta Crystallogr D Struct Biol 76, 724–728 (2020).

47. Tan, Y. Z. et al. Addressing preferred specimen orientation in single-particle cryo-EM through tilting. Nat Methods 14, 793–796 (2017).

48. Pettersen, E. F. et al. UCSF Chimera--a visualization system for exploratory research and analysis. J Comput Chem 25, 1605–1612 (2004).

49. Emsley, P., Lohkamp, B., Scott, W. G. & Cowtan, K. Features and development of Coot. Acta Crystallogr D Biol Crystallogr 66, 486–501 (2010).

50. Lebedev, A. A. et al. JLigand: a graphical tool for the CCP4 template-restraint library. Acta Crystallogr D Biol Crystallogr 68, 431–440 (2012).

51. Yamashita, K., Palmer, C. M., Burnley, T. & Murshudov, G. N. Cryo-EM single-particle structure refinement and map calculation using Servalcat. Acta Crystallogr D Struct Biol 77, 1282–1291 (2021).

52. Liebschner, D. et al. Macromolecular structure determination using X-rays, neutrons and electrons: recent developments in Phenix. Acta Crystallogr D Struct Biol 75, 861–877 (2019).

53. Afonine, P. V. et al. Real-space refinement in PHENIX for cryo-EM and crystallography. Acta Crystallogr D Struct Biol 74, 531–544 (2018).

54. Murshudov, G. N. et al. REFMAC5 for the refinement of macromolecular crystal structures. Acta Crystallogr D Biol Crystallogr 67, 355–367 (2011).

55. Nicholls, R. A., Tykac, M., Kovalevskiy, O. & Murshudov, G. N. Current approaches for the fitting and refinement of atomic models into cryo-EM maps using CCP-EM. Acta Crystallogr D Struct Biol 74, 492–505 (2018).

